# Hippocampal engrams configure prefrontal context representations to guide flexible decisions

**DOI:** 10.64898/2026.07.06.732916

**Authors:** Joshua B. Julian, Jesse C. Kaminsky, David W. Tank, Carlos Brody

## Abstract

Flexible behavior requires using past experiences to configure cortical computations to suit current task demands. A central question in neuroscience is how memory representations control such reconfigurations. Although hippocampal (HPC) engrams can drive learned behaviors, prior studies have been largely limited to a fixed stimulus-response computation. Thus, whether engrams can drive retrieval of a set of stimulus-response mappings, rather than a specific response itself, remains unresolved. Moreover, how engrams affect cortical task representations and dynamics so as to produce engram-consistent behavior remains unstudied. Here, we address both questions by combining tagging and reactivation of HPC engrams with simultaneous large-scale recordings in medial prefrontal cortex (mPFC) during a context-dependent task-switching paradigm in mice. We report that HPC engram reactivation caused mice to apply engram-consistent decision-rules rather than a specific motor output. Simultaneous mPFC recordings revealed that reactivating the HPC engram for a given context reinstated the representation of that context in mPFC within hundreds of milliseconds, indicating that it was mediated by rapid network effects, leading to choice behavior consistent with the reactivated HPC engram. Other aspects of endogenous dynamics in mPFC were left remarkably intact. Together, our findings provide direct causal evidence that HPC engrams can configure task-relevant population states of downstream cortical circuits in real time, establishing a neural mechanism by which memory traces control flexible behavior.

## Main Text

Flexible behavior depends on using context to select the appropriate set of actions that lead towards achieving one’s goals. This requires each context to engage a distinct configuration of cognitive processes, or ‘task set’, often operationalized in task-switching studies as the context-specific rules linking sensory stimuli with task-appropriate actions (Monsell 2003; Sakai 2008). Network models of task-switching typically represent context as a scalar input of unspecified biological origin (Dubreuil et al. 2022; Flesch et al. 2022; Pagan et al. 2025; Langdon and Engel 2025). Nevertheless, the hippocampus (HPC) has long been implicated in encoding context across many domains (Maren et al. 2013; Hirsh 1974; Smith and Mizumori 2006; Epstein et al. 2017; Cohen and Eichenbaum 1993; O’Keefe and Nadel 1978), positioning it as a plausible source of contextual signals that specify task sets during flexible behavior.

Activity-dependent tagging provides a powerful approach to test the causal role of context-dependent HPC representations. It enables selective labeling and manipulation of HPC neurons active during contextual learning (referred to as “engrams”) (Josselyn and Tonegawa 2020). Foundational studies demonstrated that artificially reactivating HPC engrams can drive context-dependent behaviors (Garner et al. 2012; Liu et al. 2012; Ramirez et al. 2013; Coelho et al. 2024; Redondo et al. 2014). Yet to our knowledge, these demonstrations have all been performed within a single task set. For example, in a typical context-dependent place preference engram study the rule linking sensory stimulus and action is fixed throughout the experiment (“sensory stimulus indicating context A → go to place X”). A similar description applies to fear conditioning and place avoidance engram studies. Whether or not HPC engrams can causally drive task sets thus remains unresolved, with important implications for both the engram and cognitive control literatures.

Moreover, HPC neurons are not themselves motoneurons. To affect behavior, HPC engram reactivations must influence either downstream motor representations themselves, or the computations that lead to motor activity. How this occurs, and the specific pathways involved, also remains unknown. Activity-dependent fluorescent labeling across the brain during HPC engram reactivation has implicated a broad spectrum of brain regions as potential downstream targets of the reactivation (Roy et al. 2022; Dorst et al. 2024; Tanaka et al. 2014; Guskjolen et al. 2018). But to our knowledge, no study has yet simultaneously reactivated HPC engrams and recorded downstream cortical activity during behavior, leaving the downstream neural dynamics by which engrams drive context-dependent behavior largely uncharacterized. This gap is especially critical because engram reactivation studies typically impose artificially synchronous activity (e.g., optogenetically), potentially obscuring how engrams operate under natural conditions and how their activity is translated into behaviorally-relevant downstream signals.

Existing data thus leave several critical questions entirely open (**Fig. 1A**). First, are HPC engram reactivations restricted to driving context-dependent motor actions? Or are they capable of driving context-dependent task sets, which are fundamentally more abstract? Second, do behaviorally-relevant downstream effects occur on a timescale of seconds, consistent with synaptic activations? Or over minutes, consistent with slower neuromodulatory processes? Third, can they drive explicit representations of context in downstream regions? Or do they influence downstream computations without modifying existing context representations? And fourth, does the artificially synchronous HPC stimulation used in most engram reactivation studies lead to downstream activity that preserves core features of natural spatiotemporal circuit dynamics? Or are the downstream natural dynamics fundamentally disrupted?

**Figure 1.**
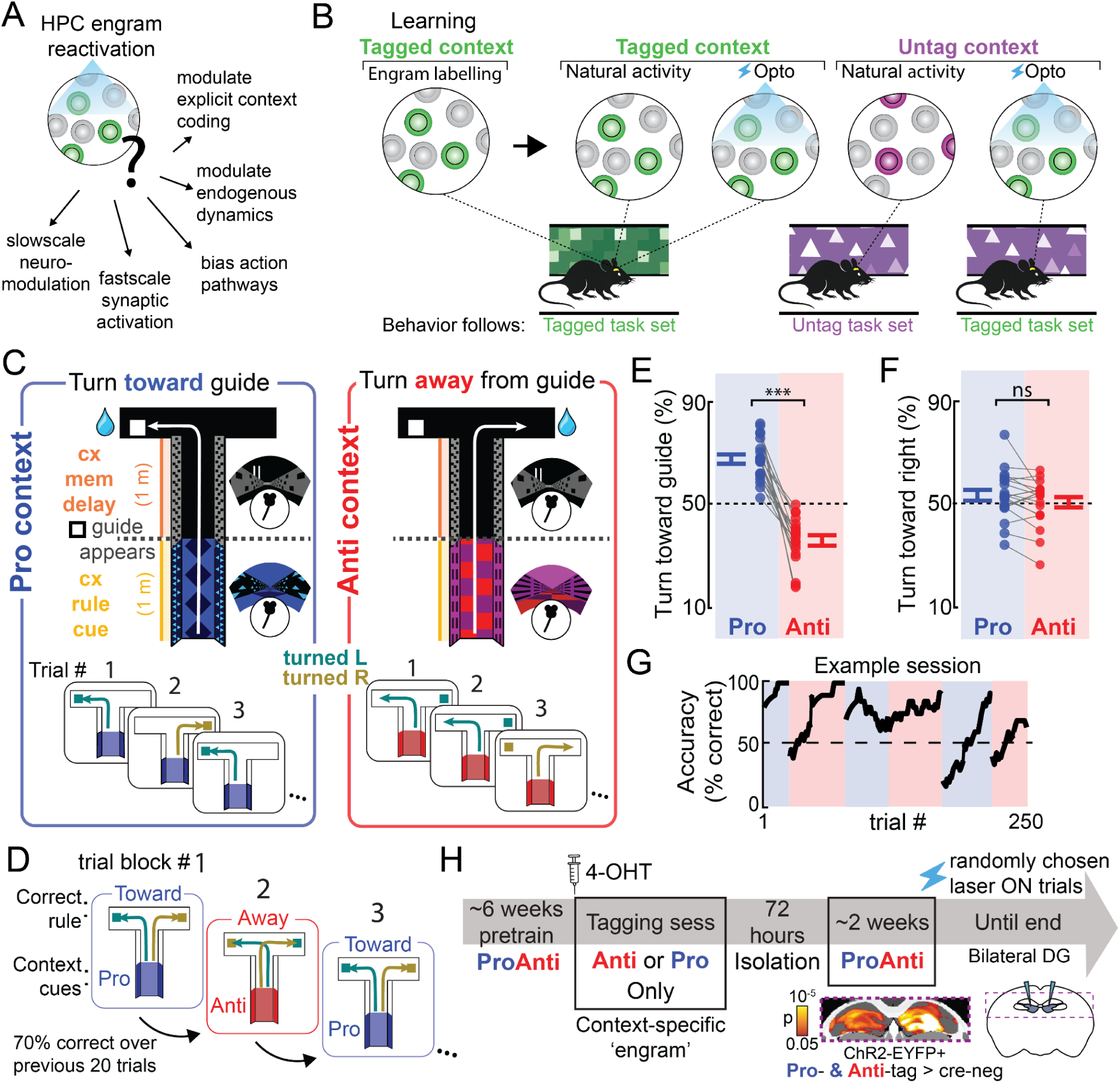
Pro-Anti task-switching paradigm and predictions. **(A)** Possible downstream mechanisms by which HPC engrams drive context-dependent behavior. **(B)** During context-dependent decision-making, the same HPC ensemble is predicted to be active during laser-off and opto trials in the tagged context, producing behavior that follows that the tagged task set. In the untagged context, different ensembles are predicted to be active during laser-off and opto trials, causing behavior to follow the tagged task set only during opto engram reactivation. **(C)** Mice performed a behavior involving different sensorimotor rules in different contexts: in the Pro context (blue), the rule was to turn toward the visual turn guide; in the Anti context (red), turn away from it. The context rule cue was presented only in the first half of the stem, followed by the context memory delay during which the guide appeared. In the delay the two contexts were identical to each other, requiring contextual identity to be held in memory from that point on. Guide position (left vs. right side of the T-maze) was balanced across trials. Cyan and gold refer to separate left and right choice trials, respectively. **(D)** Contexts were presented in interleaved blocks, and switched once performance exceeded 70% correct over the previous 20 trials. **(E)** Decision rule by context (probability of turning toward the guide) during pre-tagging sessions. Mice turned toward in Pro and away in Anti (N=20; ***p=2x10^-6^). **(F)** Motor choice (probability of right turn) was similar across contexts (p=0.26). **(G)** Example session showing performance (% correct, 10-trial within-block moving mean). **(H)** Experimental timeline. Bottom: significance map of ChR2–EYFP expression (Pro & Anti-tag > Cre- controls; N = 12 vs. 4) in HPC, overlaid on atlas (p < 0.05, SVC). In (E-F), dots represent individual mice; error bars denote mean ± SEM.

Here we set out to answer these questions with a combination of a novel mouse task-switching behavior, HPC engram reactivation, and simultaneous electrophysiological recordings in one potential downstream target, the medial prefrontal cortex (mPFC) (Thierry et al. 2000; Hoover and Vertes 2007; Ye et al. 2017; Eichenbaum 2017). We chose the mPFC as a strong candidate for observing downstream effects of HPC engram reactivation because decades of research have established that the PFC is required for task set updating (Durstewitz et al. 2010; Birrell and Brown 2000; Milner 1963; Miller and Cohen 2001), and coordinated HPC–mPFC activity is critical for memory-guided planning and decision-making more broadly (Wikenheiser et al. 2017; Weilbächer and Gluth 2016; Tavares and Tort 2022; Tang et al. 2021; Elston and Wallis 2025; Jadhav et al. 2016; Eichenbaum 2017; Spellman et al. 2015; Place et al. 2016).

Our results show that HPC engrams are capable of driving downstream task sets, without directly driving motor representations; they do so within hundreds of milliseconds of reactivation, thus establishing a key role for synaptic and network effects in the engram→behavior pathway; they modify existing context representations; and despite the artificial nature of laser stimulation, they preserve major downstream spatiotemporal dynamics features, suggesting that the artificial aspects of the reactivation are filtered by the network into downstream activity that closely mimics natural activity.

## A mouse task-switching behavior

We first set out to answer whether HPC engrams can causally drive task sets (**Fig. 1B**). To dissociate motor action from task set specification, we designed a task-switching behavior in which context determines which sensorimotor decision rule, rather than motor action, to apply on each trial. Following task-switching paradigms in non-human primates (Munoz and Everling 2004; Mante et al. 2013; Wu et al. 2020; Elston and Wallis 2025; Siegel et al. 2015; Bernardi et al. 2020), and their adaptation to rodents (Pagan et al. 2025; Duan et al. 2015, 2021; Wimmer et al. 2015), we developed a behavioral task for head-fixed mice in virtual reality (VR) based on the classic “Pro/Anti task-switching” paradigm, in which context determines whether subjects orient toward or away from a stimulus (**Fig. 1C**).

Mice ran up the stem of a VR T-maze, and were trained to make a decision as to whether to turn left or right at the top of the “T”. The T-maze could be rendered in one of two contexts (labelled “Pro” and “Anti” in **Fig. 1C**), each of which was associated with a different sensorimotor rule. The contexts were distinguished by salient multisensory cues presented exclusively along the first half of the T-maze stem (the “rule cue” zone), and were presented in interleaved blocks of trials within each experimental session (**Fig. 1D**). Upon exiting the rule cue zone and entering the second half of the stem (referred to as the “contextual memory delay” zone), a salient visual stimulus called a turn “guide” was presented either on the left end of the VR corridor or on the right end (balanced across trials). The guide was visible throughout the memory delay zone of the T-maze stem. In the Pro context, the task-appropriate sensorimotor rule for receiving water reward was to turn *toward* the guide at the top of the stem (regardless of whether the guide was on the left side or right side). In the Anti context, the task-appropriate sensorimotor rule was to turn *away* from the guide. The memory of the context thus did not specify a particular “go-left” or “go-right” motor action, but instead specified the rule to apply.

Once trained, mice showed a strong effect of context on their decisions (**Fig. 1E**; for training procedure, see **Fig. S1**). Consistent with the task design, there was a negligible effect of context on their overall rightwards versus leftwards choices (**Fig. 1F**). Within each session, mice ran a median (± 1MAD) of 280±37 trials, and completed 4±1 context blocks (**Fig. 1G**; **Fig. S2A**). Control manipulations of the contextual cues confirmed that these multisensory cues played a role in how mice identified context and retrieved the associated decision rule, rather than exclusively inferring the current decision rule from recent trial outcome feedback (**Fig. S2E-G**). Switching contexts across blocks incurred a transient performance cost, consistent with perseveration of the previous context’s decision rule (**Fig. 1G**; **Fig. S2B**). Asymmetric task switch costs are often found in other species (Monsell et al. 2000; Allport et al. 1994; Duan et al. 2015); here the switch cost was greater when switching from Pro to Anti than vice versa **(Fig. S2C-D).**

## HPC engrams drive context-dependent sensorimotor task sets

To test whether context engrams contribute to this task-switching behavior, we used activity-dependent optogenetics to label and later reactivate HPC ensembles associated with either the Pro or the Anti context. We crossed mice expressing tamoxifen-dependent CreERT2 under the *Arc* promoter with a Cre-dependent ChR2-EYFP reporter line, so that neurons active shortly following tamoxifen injection permanently express ChR2 (Denny et al. 2014; Guenthner et al. 2013). To target context-specific ensembles and avoid environmental novelty-related ensembles (Cleland et al. 2017), mice were first pretrained on the ProAnti task (**Fig. S1A**). They were then injected with 4-hydroxytamoxifen (4-OHT) and completed one training session that consisted of trials of only a single context (**Fig. 1H**). In the “Pro-tag” group of mice, this session had only Pro context trials; In the “Anti-tag” group, this session had only Anti context trials. This single-context training session labeled neurons active in their specific tagging context with ChR2-EYFP (**Fig. S3A**).

Behavioral performance improved during the tagging session (**Fig. S1B**), confirming that animals continued to strengthen their association between a context and its corresponding decision rule. Following a three-day isolation period, mice were retrained to criterion. In subsequent sessions, the tagged population was reactivated during select task epochs by delivering blue laser light (∼2 mW, 20 Hz) through bilateral optic fibers targeting the dorsal dentate gyrus (DG; **Fig. S1C**, **S3B**). This was done on 30% of randomly selected trials, with the remaining interleaved 70% of trials serving as laser-off controls. To describe both Pro-tag and Anti-tag groups of mice together, we will refer to the context matching an animal’s tagging context as the “tagged” context, and the other context as the “untagged” context.

If HPC engrams contribute to flexible decision-making, reactivating context-specific ensembles should retrieve the associated decision rule and bias choices accordingly. During tagged context trials, behavior during laser-on trials could be similar to behavior during laser-off (control) trials, particularly if HPC activity was already correctly representing the tagged context before the laser is turned on. In contrast, during untagged context trials, turning the laser-on should cause mice to flip from the untagged to the tagged representation, and disproportionately apply the tagged rule. We thus expect the most salient effects to be seen in the untagged context. Overall, the pattern of predicted effects is depicted in **Fig. 2A**.

**Figure 2.**
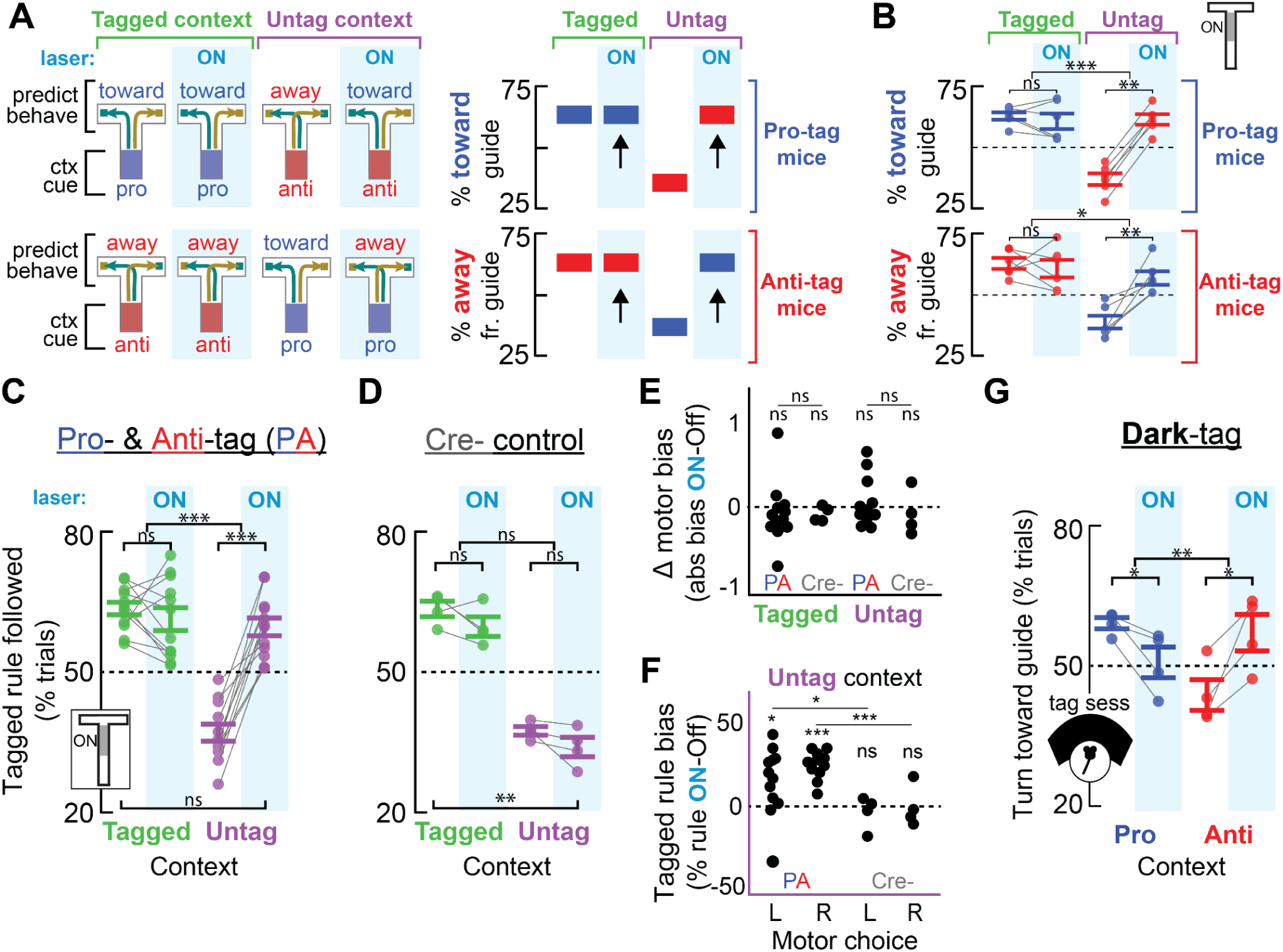
Context-specific hippocampal engrams bias context-dependent decisions. **(A)** Predicted behavioral effect: reactivating tagged engrams should bias mice to use the tagged choice rule in both Tagged and Untagged contexts, in both Pro- (top row) and Anti- (bottom row) tagged mice. Behavior is thus predicted to be tag-consistent (black arrows) independent of the experimenter cued context. **(B)** Observed: memory delay laser stimulation increased use of the tagged rule in the Untagged context, but there was no effect of the laser in the Tagged context, in both Pro-tag (top, N=6) and Anti-tag (bottom, N=6) mice. **(C)** Combining Pro- and Anti-tag mice (PA, N=12), memory delay stimulation increased use of the tagged rule in the untagged context (above Off, p=0.000001, and above 50% chance, p=0.0004). In the tagged context, PA used the tagged rule during both laser-off and -on trials, with no additional effect of the laser (context × laser interaction: p=0.00002). **(D)** Cre- controls (N=4) showed no laser effects in either context. **(E)** No changes in absolute motor-choice bias within or across groups in either context. **(F)** Tagged rule bias in the Untag context in PA present during both left and right choice trials (both p<0.02), absent in Cre-. **(G)** Dark-tag controls (N=4), tagged in darkness without a task, were generally impaired by laser stimulation (context x laser interaction: p=0.007). Dots denote individual mice; error bars denote mean ± SEM. *p<0.05, **p<0.01, ***p<0.001. Full statistics in Table S1.

**Fig. 2B-C** shows the experimentally obtained effects when context-specific HPC ensembles were optically stimulated during the contextual memory delay (i.e., the period during which mice must maintain context in memory to guide their choice; effects of rule cue zone reactivation are described below). The observed effects were remarkably similar to the predictions (compare **Figs. 2A and 2B**). Strikingly, choices on laser-on trials in the untagged context were indistinguishable from choices on laser-off trials in the tagged context, indicating that artificial reactivation substituted for the natural contextual cues. We found no significant effect of reactivation in the tagged context, suggesting that the strength of an already-represented tagged context was not boosted further by optical reactivation. In sum, reactivation of HPC engrams in the memory delay zone drove tag-consistent flexible decision-making: turning the laser-on led mice to behave as if they were in the tagged context, even when tested in the opposite context.

To confirm that the tag-specific decision bias was not an artifact of optical stimulation alone, we repeated the experiment in control mice that underwent the same procedures but lacked the Cre allele (Cre- group). In these Cre- mice, optical stimulation had no effect (**Fig. 2D**), ruling out nonspecific stimulation effects and demonstrating that the behavioral bias depended on tagged HPC ensembles.

The context-specific bias induced by HPC memory delay zone reactivation was independent of which context was tagged, with similar effects in both the Pro- and Anti-tag groups (**Fig. 2B**; **Fig. S4A-B**). The tag-specific decision bias was robust across sessions (observed in 87.8% of sessions; sign test, p=1.37×10^-8^; **Fig. S4C–D**) and across trials within a block, with reactivation causing tag-consistent choices to occur more frequently in the untagged context regardless of time since context switching (**Fig. S4E**). Control analyses confirmed that this effect was not due to nonspecific changes in VR locomotion (**Fig. S5**).

To test whether reactivation of context-specific HPC cells drives decision rule retrieval rather than simply biasing motor output, we first confirmed that optical stimulation did not alter overall motor choice bias (**Fig. 2E**). We then calculated the decision rule bias separately for left-and right-turn choice trials. Reactivation of context-specific HPC ensembles caused mice to apply the tagged rule in the untagged context for both turn directions (**Fig. 2F**). These results show that HPC engram reactivation influences retrieval of the decision rule itself, rather than producing a motor bias.

If HPC engrams provide a neural source of context during flexible decision-making, activating HPC cells unrelated to a task-relevant context should disrupt performance. To test this, we applied the same experimental procedure to a separate group of mice, except that following 4-OHT treatment to induce tagging, these mice were head-fixed in the VR rig in the dark and received randomly interspersed rewards (Dark-tag group; **Fig. S6A–D**). For this Dark-tag group, memory delay zone reactivation produced choices that were at chance in both contexts (**Fig. 2G**), a markedly different effect compared to the Pro- or Anti-tagged mice. Overall performance (% correct) during laser-off trials was also reduced in Dark-tag mice relative to pre-perturbation sessions (**Fig. S6E**). In a separate group of VGAT-ChR2 mice, optogenetic inhibition broadly targeting the DG also generally impaired ProAnti behavior (**Fig. S6F**). These results indicate that the context-specific decision bias in Pro- and Anti-tag mice requires activation of a HPC ensemble associated with the tagging context, and that stimulating a HPC population unrelated to decision-making interferes with general task performance rather than eliciting context-specific decisions.

The preceding results involved reactivation of context-specific HPC cells during the memory delay zone of the stem of the T-maze, a period during which mice must maintain contextual information in memory to guide their choices. To test whether HPC engrams also contribute during other task epochs, we also reactivated HPC engrams in Pro- and Anti-tag mice during the whole stem of the VR T-maze (i.e., both the rule cue and memory delay zones), or during the rule cue zone alone. Whole-stem reactivation biased choices toward the tagged rule similar to memory delay reactivation alone (**Fig. S7A**). But in contrast, reactivation during the rule cue zone failed to produce a tag-specific bias: behavior on laser-on trials was impaired in both contexts, with the strongest deficit in the Anti context regardless of tag identity (**Fig. S7B–C**). Rule cue-only reactivation also decreased mouse running speed specifically at the transition between the rule cue zone and the memory delay (**Fig. S5B**). Memory delay stimulation produced a significantly stronger tag-specific bias than rule cue–only reactivation (p=0.0002; **Fig. S7D**), indicating that HPC engram activity is particularly critical for decision-making when contextual information must be internally maintained and a context-dependent choice is made.

## HPC engrams drive mPFC contextual representations

Building on these behavioral findings, we next sought to determine how HPC engram reactivation shapes downstream neural computations that implement task sets. We chronically recorded single-unit activity in the mPFC during simultaneous ProAnti task performance and HPC engram reactivation (**Fig. 3A**), allowing us to directly examine how context-specific HPC output influences prefrontal representations supporting flexible behavior. Recordings targeted anterior cingulate and prelimbic cortices (296 ± 82 simultaneously recorded units, median ± s.d., per behavioral session; **Fig. S3C**).

**Figure 3.**
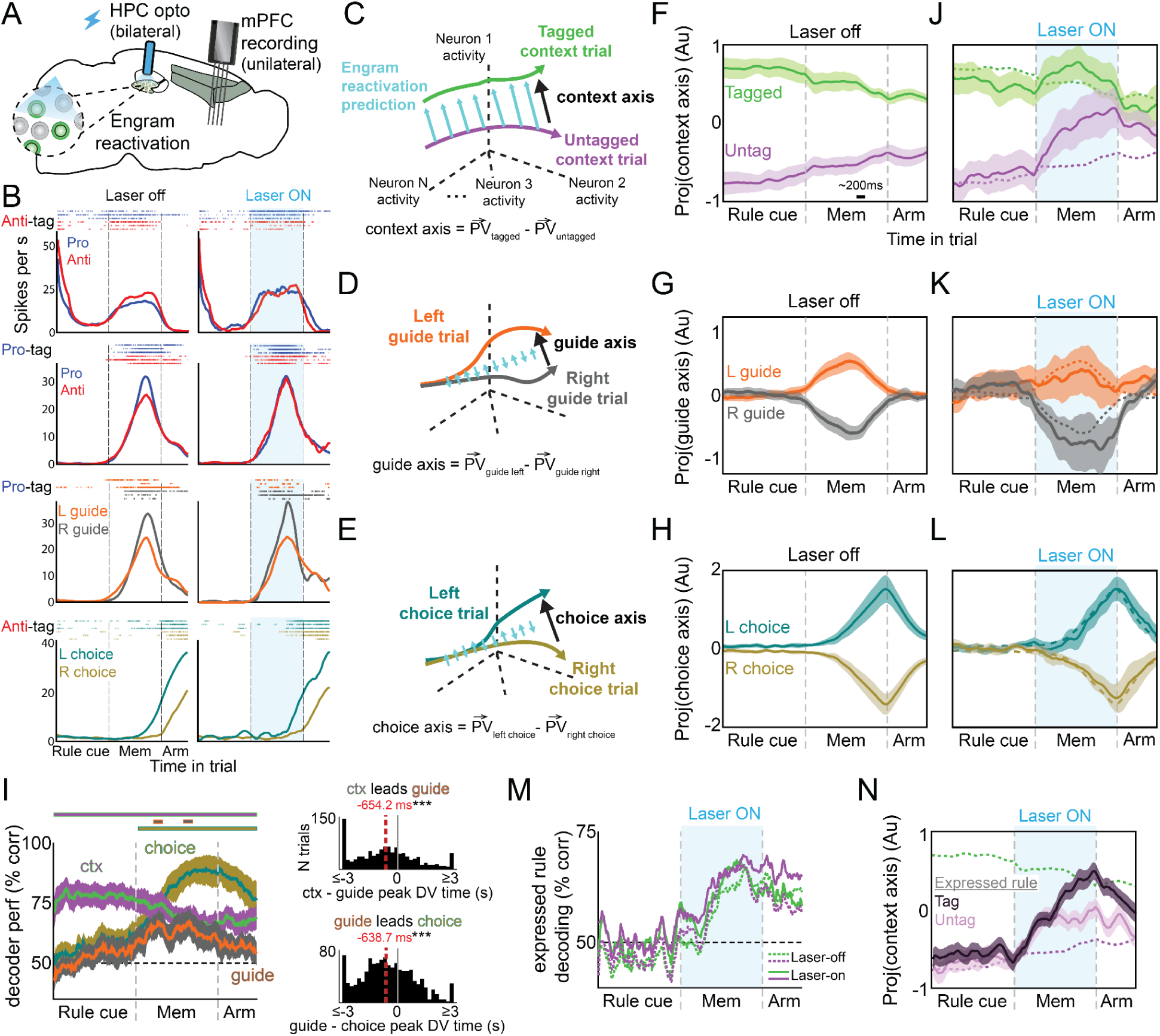
HPC engram reactivation reinstates tagged context representation in mPFC. **(A)** mPFC single-unit activity recorded using Neuropixel probes during simultaneous ProAnti behavior and HPC engram reactivation (N=2 Pro-tag, N=2 Anti-tag). **(B)** Example mPFC single-unit activity during laser-off (left) and memory-delay reactivation (right) trials for four different units (rasters from 6 random trials). **(C-E)** Schematics of context, turn guide side, and choice coding axes, and predicted effects of engram reactivation. (**F-H**) Population trajectories over time projected onto the (F) context axis, (G) guide axis, (H) and choice axis during correct laser-off trials (N=11 sessions). Scale bar in (F) indicates elapsed time during the memory delay. **(I)** Left: Trialwise LDA decoding of context, turn guide side, and choice over time during laser-off trials. Significance markers show above-chance (50%) decoding (p<0.001). Right: on individual trials, the time of the absolute peak of the context decision-variable (DV) preceded guide DV (top: sign-test, p=3.25x10^-30^) and guide preceded choice DV (bottom: p=5.86x10^-26^). (**J-L**) Projections onto the (J) context, (K) guide, (L) and choice axes during memory delay reactivation trials. Dotted lines denote respective laser-off baselines. **(M)** LDA decoding of expressed decision rule (Pro vs. Anti) during laser-off (dotted) and memory delay reactivation (laser-on; solid) trials, shown separately by context. Decoders were trained on laser-off trials controlling for contextual cues, motor choice, and performance. No significant differences between laser-on and -off trials in either context, indicating that the laser induced shift in expressed rule was represented in mPFC. Curves show bootstrap mean; overlapping error bars omitted for clarity. **(N)** Same as (J) in the untagged context, but split by expressed behavioral rule (mean ± SEM pooled across trials; statistical inference via bootstrap). (F-L) shaded lines denote bootstrap mean ± SD. Full statistics in Figs. S10-S12, and Table S1.

Consistent with previous studies involving rodent navigation (Kaefer et al. 2020; Hok et al. 2005; Zielinski et al. 2019; Sauer et al. 2022), we found that mPFC had substantial place-field-like activity, with most neurons having a preferred location of space in which they tended to have high activity. As the animal traverses the maze, this led to an orderly sequence of activated neurons that tiled the T-maze (example neurons **Fig. 3B** and **Fig. S8A**; see **Fig. 4** for further characterization of these sequential dynamics). Successful performance of the ProAnti task requires animals to integrate contextual information with turn-guide side to select the appropriate motor response on each trial. Consistent with nonhuman primate and rat studies (Rigotti et al. 2013; Mante et al. 2013; Euston et al. 2012; Spellman et al. 2015; Duan et al. 2021), during laser-off trials, individual mouse mPFC neurons encoded these task variables, including context, turn-guide side, and upcoming choice. The variables were encoded as modulation of the underlying place-dependent activity (**Fig. 3B**; **S8A-B**).

**Figure 4.**
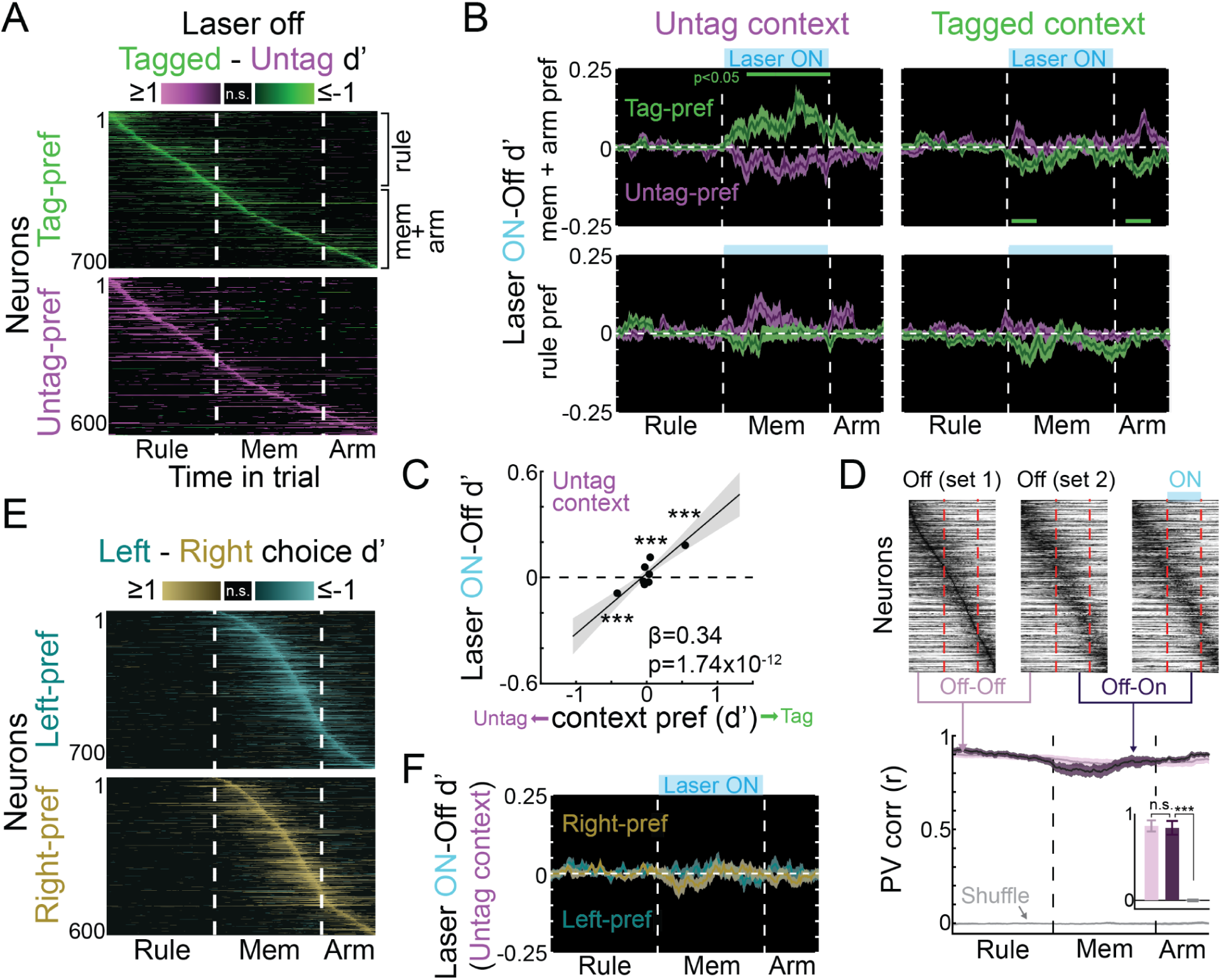
Single-unit contributions to HPC-driven context reinstatement in mPFC. **(A)** Context discriminability (d′) in context modulated cells during laser-off trials (inclusion threshold: p<0.01, uncorrected). Neurons are sorted by peak discriminability time and context preference. **(B)** Laser modulation (on–off d′) for neurons grouped by rule cue or memory and arm field location, and context preference, as in (A), shown separately in untag and tagged contexts (median ± bootstrap SEM). During engram stimulation in the untagged context, mem + arm tag-preferring neurons were excited (p<0.05, FWE-corrected), whereas mem + arm untag-preferring neurons showed weaker, non-significant suppression. Reactivation in the tagged context produced brief suppression of tag-preferring neurons at laser onset and offset. Rule cue neurons were not significantly modulated. **(C)** Mean laser modulation vs. laser-off context discriminability during the 2nd half of the memory delay, pooled across neurons (untag context; n=1992). Laser effects scaled with context tuning: untag-preferring neurons were significantly suppressed and tag-preferring neurons excited (linear mixed model, shading denotes 95% CI). Points show means of eight equally sized context preference bins (sign-rank, FDR-corrected: bin #1 p=4.9x10^-5^, 2-6: >0.05, 7: 0.0007, 8: 1.1x10^-8^). **(D)** Top: Untag context activity traces sorted by peak firing time during an example laser-on–trial-matched subset of laser-off trials (set 1). Rows are max-normalized per condition. Bottom: time-resolved population vector (PV) correlation across conditions (mean ± bootstrap SEM). Shuffle permutes neuron identities in laser-off activity for the laser-off vs. -on comparison. Inset shows average similarity over the memory delay (±1 SD; t-tests: off vs. on: p=0.35, on vs. shuffle: 9.9x10^-12^). **(E)** Same as (A) split by choice preference. **(F)** No laser modulation in choice-preferring neurons during stimulation in the untag context. *p<0.05, **p<0.01, ***p<0.001.

To characterize how these key task variables were represented at the population level, we analyzed laser-off activity in neural state space, with each dimension corresponding to a single neuron. We identified the coding directions in this space that maximally discriminated context, turn guide side, and upcoming choice (Inagaki et al. 2019; Li et al. 2016) (**Fig. 3C-E**). These coding directions were largely orthogonal to each other, indicating that each of these task variables was represented along distinct axes of population activity (**Fig. S9**).

Projecting single-trial population activity onto the context axis revealed well-separated context-specific distributions throughout the entirety of laser-off trials (**Fig. 3F**; **S10A**). In contrast, projections onto the guide and choice axes diverged following the appearance of the turn guide (**Figs. 3G-H**; **S11A; S12A**). Linear discriminant analysis (LDA) revealed a temporal ordering to the emergence of these representations on individual trials: consistent with subjects using the Pro/Anti trial block structure, information about context was present from the start of each trial; upon the start of the memory period/presentation of the turn-guide, a representation of turn-guide side appeared, followed by a representation of motor choice (**Fig. 3I**). Thus, mPFC population activity mirrors the sequence of computational task demands, representing context, turn guide, and motor choice in the order expected for accurate performance. We also asked whether mPFC activity encoded the decision rule that the animals expressed in their behavior. Because the animals made errors, we defined the expressed decision rule on each trial as Pro if the animal turned towards the guide, and as Anti if the animal turned away from it, independent of the rule cued by the experimenters. LDA found reliable decoding of the expressed decision rule in laser-off trials, even when controlling for contextual cues, motor choice, and balancing the number of correct and incorrect trials used in the analysis (**Fig 3M**). Thus, during control trials, both the cued decision rule and the expressed decision rule are represented in mPFC during the memory delay, with the latter ramping up during the memory delay.

Having established that mPFC encodes the variables required to solve the ProAnti task, we next asked whether and how these representations are causally influenced by HPC engram reactivation. If HPC engrams bias motor action pathways, we would expect perturbations along the choice axis. Alternatively, if HPC reactivation conveys contextual information that specifies the active task set, its primary effect should be selective to the context axis. In this latter framework, HPC reactivation would influence behavior by altering the contextual state from which choice computations proceed, rather than by directly modifying motor choice representations themselves. Since choice representations emerged following context, activity along the choice axis itself may thus remain largely unperturbed despite reliable changes in choice behavior.

mPFC neurons were not directly activated by HPC laser stimulation (**Fig. S13**), indicating that any observed effects reflected network-level modulation, likely through polysynaptic pathways (Eichenbaum 2017). During memory-delay stimulation, HPC engrams selectively altered activity along the context axis: In the untagged context (purple traces in **Fig. 3I**), reactivation shifted mPFC population activity along the context axis toward the tagged-context trajectory, whereas no significant effect of the laser was observed in the tagged context (green in **Fig. 3I**). By the end of the memory delay in laser-on trials, tagged and untag context trajectories became statistically indistinguishable (**Fig. S10A**). Across trials, HPC-driven contextual reinstatement in mPFC occurred at variable times throughout the memory delay, with a median shift time of ∼850 ms following laser onset (**Fig. S14A**). Importantly, the context axis accounted for a comparable fraction of population variance during laser-off and laser-on trials (**Fig. S10B**), demonstrating that engram reactivation did not change the contribution of context coding but instead biased activity within this subspace. Thus, memory delay reactivation reinstated an explicit engram-associated context representation in mPFC.

In contrast, trajectories projected onto the guide and choice axes remained well separated during memory-delay HPC reactivation, with no significant effect of laser stimulation on how these variables are represented, either overall (**Fig. 3J-K**; **S11A; S12A**) or when analyzed separately in each context (**Fig. S11B-C; S12B-C**). In other words, activity along the choice axis aligned with the animal’s expressed choice in the untagged context, even when HPC reactivation caused a change in expressed rule (**Fig. S12D**). Notably, on trials where HPC reactivation caused mPFC contextual reinstatement, robust choice representations emerged following that reinstatement (**Fig. S14B**), suggesting that laser effects on choice are mediated specifically through effects on the represented context. Guide representations were unrelated to the timing of contextual reinstatement (**Fig. S14C**). These results are consistent with the idea that an HPC-driven contextual state in mPFC gates the emergence of downstream choice representations.

Consistent with the nonspecific behavioral effects of rule-cue-period-only stimulation, ambiguous neural effects were observed in mPFC context representations for trials with engram reactivation during this epoch. This was particularly salient in the Anti context regardless of tag identity (**Fig. S15A-D**). As during memory-delay-period HPC stimulation, rule cue reactivation had no significant effect on the guide and choice dimensions (**Fig. S15E-F**).

If HPC engrams establish the contextual input for flexible decision-making, then reinstating context in mPFC should place the network into a state consistent with its behavior on that trial. To test this, we asked whether the expressed rule decoder, trained on laser-off trials, also decoded the expressed rule during laser-on trials. We found that this decoding was intact: as with guide and choice, HPC engram reactivation had no significant effect on how the expressed rule was represented (**Fig 3M**). To ask what distinguished trials in which reactivation did or did not induce a change in expressed rule, for each of these two groups of trials we examined the time-course of the context representation during laser-on trials in which the cued context was the untagged context. We found that trajectories along the context axis initially shifted toward the tagged context representation regardless of the expressed rule. Yet, this shift only persisted on trials in which the tagged rule was ultimately expressed (i.e., when cued and expressed rules did not match, **Fig. 3N**; **Fig. S14D**). When reactivation failed to induce the tagged decision rule (i.e., when cued and expressed rules matched), population activity returned to the untagged trajectory by the end of the trial. These results indicate that HPC engrams transiently reinstate the tagged context in mPFC independently of behavior, but that only sufficiently sustained contextual bias leads the network to stabilize in a tag-rule-consistent state. These data demonstrate that HPC engrams modulate mPFC computations on a timescale of hundreds of milliseconds. To provide an independent, mechanistically-direct readout of the timing of HPC→mPFC transmission, we also examined the entrainment of mPFC spiking to the 20-Hz HPC stimulation frequency. Fourier analysis of single-neuron spike trains revealed a robust 20-Hz peak during laser-on relative to laser-off trials, demonstrating frequency-specific entrainment of mPFC to HPC-driven activity (**Fig. S16A-C**). Time-resolved wavelet analysis further showed that engram reactivation elicited a rapid, broadband increase in mPFC spiking ∼100 ms after laser onset, likely reflecting a transient increase in mPFC excitability, followed by disproportionate entrainment of mPFC spiking to the 20-Hz stimulation frequency emerging ∼800 ms after laser onset (**Fig. S16D-F**). Together, the contextual representation and entrainment results converge on the conclusion that HPC engrams drive mPFC contextual reinstatement through multisynaptic transmission and rapid (hundreds of milliseconds) network effects.

The HPC-driven population-level contextual reinstatement observed in mPFC could, in principle, arise through excitation of neurons preferring the tagged context, suppression of neurons that prefer the untagged context, or a combination of both. It could also reflect changes in a small set of highly responsive neurons or distributed modulation across a broader context-tuned ensemble. To distinguish these possibilities, we examined the contribution of context to single-neuron mPFC activity (**Fig. 4A**). During memory-delay reactivation in the untagged context, neurons preferring the tagged context and with a firing field in the memory delay zone or arm of the T-maze exhibited a robust increase in firing rates (**Fig. 4B**, left upper). Conversely, neurons preferring the untagged context showed firing rate suppression, albeit with weaker modulation than tagged-preferring cells. No effect of memory delay zone reactivation was seen in cells with firing fields in the earlier rule zone (**Fig. 4B**, left lower). As a result, population activity during stimulation shifted toward a pattern resembling that observed in the tagged context during laser-off trials. This shift reflected a coordinated, population-wide modulation that scaled with each neuron’s degree of context tuning, rather than being driven by a small subset of highly responsive units (**Fig. 4C**).

HPC reactivation in the tagged context produced only brief, weak suppression of tag-context-preferring neurons following laser onset and offset without sustained population-wide modulation (**Fig. 4B**, right). This is consistent with the absence of HPC stimulation effects on mPFC population activity during tagged context trials (**Fig. 3J**). Neuronal activity was also not modulated as a function of their choice tuning (**Fig. 4E, F**). Finally, reactivation in the rule cue zone failed to elicit tag-consistent modulation, instead biasing activity toward a more Pro-context-like pattern in the Anti context regardless of tag identity, paralleling the observed change in behavior (**Fig. S15G**).

The lack of an effect of memory-delay HPC reactivation on mPFC neurons with firing fields in the rule cue zone suggests that reactivation may bias mPFC contextual representations without reorganizing spatiotemporal sequence dynamics. Indeed, direct examination of mPFC sequential activity revealed that although a small number of neurons altered their sequential firing during reactivation (**Fig. S8C**), spatiotemporal sequences were strongly preserved at the population level (**Fig. 4D**). Bayesian decoding of spatiotemporal position from mPFC activity confirmed this observation: decoders trained on laser-off trials transferred to reactivation trials with comparable accuracy, with only a small increase in decoding error comparable to the error observed when decoding position across contexts during laser-off trials (**Fig. S17A-B**). This error increase during HPC stimulation was driven by a small (∼3.4 cm) systematic backward shift in decoded position, consistent with a small spatiotemporal advance in sequential firing rather than disruption of the sequence itself (**Fig. S17C**). Rule cue reactivation had no significant effect on sequential activity (**Fig. S17D-F**). Thus, despite the artificial HPC stimulation lasting the entirety of the memory delay, and the observed 20-Hz entrainment, engram reactivation modulated the contextual component of mPFC activity without altering sequential dynamics. This suggests that the artificial aspects of the reactivation are filtered by the network into downstream activity that closely resembles natural activity.

## Discussion

In this study, we show that HPC engrams can causally drive behaviors that require abstract, flexible task sets. Rather than eliciting specific motor action plans, reactivation of context-specific HPC engrams led mice to retrieve and apply the appropriate rule for a decision, even when that rule demanded opposing actions across trials. Therefore, engram reactivation can direct the subject’s mapping between sensory inputs and actions, the defining feature of a task set. By demonstrating that HPC engrams are causal to flexible behaviors guided by abstract cognitive states, the results presented here extend prior work that showed that reactivation of sensory- or fear-associated engrams are sufficient to direct motor actions (Liu et al. 2012; Ramirez et al. 2013; Coelho et al. 2024; Redondo et al. 2014).

Previous studies combining HPC engram stimulation with immediate early gene (IEG) mapping in downstream regions have shown that HPC engram reactivation propagates activity through HPC circuits and engages distributed downstream targets (Roy et al. 2022; Dorst et al. 2024; Tanaka et al. 2014; Guskjolen et al. 2018). However, how that reactivation-driven activity relates to behavior remains unclear. With regard to mPFC, for example, (Roy et al. 2022) found that mPFC was one of the regions showing decreased neural activity due to HPC engram inhibition. But whether or not that engram-modulated activity relates to behavior or task representations, and if so, how, cannot be determined from IEG mapping alone. Only a small number of reports have used HPC engram reactivation while recording downstream neural activity during behavior (Coelho et al. 2024; Suthard et al. 2024; Rahsepar et al. 2023; Norman et al. 2025), and these focused purely within the HPC loop, for example reactivating engrams in one HPC subregion while recording in another HPC subregion. As a result, the fundamental question of how engram-driven activity shapes neocortical dynamics and population codes to support ongoing behavior has remained unexplored. Here, we show that engram reactivation selectively shifts mPFC population activity along a task-relevant axis predictive of flexible decision-making, thus linking circuit-level reactivation to cognitive function.

HPC reactivation could influence behavior either by biasing downstream motor representations directly or by reinstating an abstract contextual state that specifies the active task set. In the former case, effects should appear along motor-choice-related neural dimensions, whereas in the latter, they should be confined to context-related dimensions, with changes in choice emerging from the altered contextual state. Our results are consistent with the latter account: HPC engram reactivation selectively shifted mPFC activity toward the tagged context representation observed during control trials, effectively reinstating a tag-like contextual state. This finding demonstrates that reactivation of context-specific HPC engrams is sufficient to drive mPFC representations of context, suggesting a candidate mechanism for the context-dependent remapping previously observed in mPFC (Jung et al. 1998; Sauer et al. 2022).

In contrast, choice representations in mPFC remained essentially intact following HPC engram reactivation, in the sense that they remained aligned with the animal’s expressed choice even when HPC reactivation caused a change in expressed rule. mPFC sequential activity dynamics and guide representations were also preserved. Thus, despite the highly artificial nature of the HPC laser stimulation (15 ms pulses at 20 Hz throughout the memory zone) and despite the observed 20 Hz entrainment in mPFC activity, neural dynamics in mPFC were remarkably similar to their natural dynamics in control trials. The main effect of HPC laser stimulation was restricted to a change in context representation. Even along the mPFC context coding axis itself, HPC laser stimulation did not cause activity outside the control range, and activity variance explained by this axis remained the same on laser-on versus laser-off trials. These results suggest that the system is robust to artificial stimulation, filtering it into dynamics closely aligned with its own endogenous dynamics.

Several anatomical pathways could mediate HPC’s influence over mPFC, including polysynaptic routes through thalamic, entorhinal, and retrosplenial circuits (Eichenbaum 2017). Identifying the mechanisms by which engrams modulate downstream activity to drive behavior will be greatly facilitated by identifying the timescale of the effects. Nevertheless, most prior experiments consist of minute-scale stimulation epochs (Liu et al. 2012; Ramirez et al. 2013; Coelho et al. 2024; Redondo et al. 2014), which can limit temporal resolution when assessing how engram activity impacts downstream circuits. Here, the rapid emergence of mPFC entrainment and contextual reinstatement following HPC stimulation indicates that engram-driven effects can arise through network interactions on the timescale of hundreds of milliseconds.

Notably, engram reactivation induced a context-specific rule bias only when stimulation occurred during the contextual memory delay, not during the rule cue period when contextual information was externally available. This dissociation suggests that HPC engrams may be particularly important for maintaining or reinstating task context during internally-guided decision-making, rather than during sensory identification of context. More generally, these findings suggest that the influence of HPC engrams on cortical task representations depends on behavioral state and task demands. Identifying the circuit pathways and network conditions that gate HPC-driven task set specification will be an important direction for future work.

More broadly, these results align with long-standing indexing theories of HPC function, which propose that HPC activity patterns serve as pointers capable of reinstating distributed cortical representations (Guskjolen and Cembrowski 2023; Teyler and DiScenna 1986). In this view, HPC engrams provide the contextual input that selects among competing rule representations, enabling rapid adjustment of behavioral policies. Selective modulation of downstream representational subspaces by HPC engrams may therefore reflect a general mechanism by which memory systems gate task set configuration, with implications for decisions ranging from approach–avoidance to inference and generalization.

## Data and Code Availability

Upon publication, the datasets generated during the current study will be deposited in a public repository, and associated code will be made freely available.

## Contributions

Conceptualization: J.J.; Methodology: J.J., J.K., C.B.; Investigation: J.J., J.K.; Formal Analysis: J.J., J.K.; Visualization: J.J., J.K., C.B.; Writing – original draft: J.J., C.B.; Writing – review & editing: all authors; Funding acquisition: D.T., C.B.; Supervision: D.T., C.B.

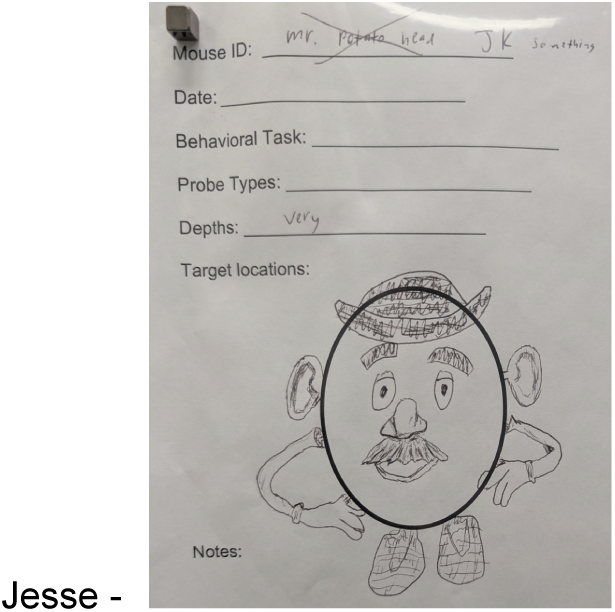

## Acknowledgements

We thank D. Allen, N. Bergevin, A. Dong, M. Hibbs, S. Janarthanan, A. Kalmbach, C. Kopec, J. LaBarbera, J. LaMonica, J. Lopez, R. Lopez, Y. Lu, A. Luna, L. Lynch, M. Magos, X. Mendez, Y. Qu, E. Rodriguez, M. Schottdorf, A. Sirko, C. Tabedzki, J. Teran, S. J. Venditto, and J. Yanar for technical assistance. We are grateful to all members of the Brody and Tank labs for their support and feedback.

This manuscript was supported in part by the National Institutes of Health (NIH) under grants 5U19NS132720 and T32MH065214. It is subject to the NIH Public Access Policy. Through acceptance of this federal funding, NIH has been given a right to make this manuscript publicly available in PubMed Central upon the Official Date of Publication, as defined by NIH. Additional support was provided by the Howard Hughes Medical Institute Investigator Program, and the Simons Collaboration on the Global Brain.

## Materials & Methods

### Animals

We used male and female mice aged 2–12 months. For our primary experiments, we used mice expressing Arc-cre-dependent ChR2, backcrossed to a C57BL/6J background (Jackson Laboratory, strain #000664) and maintained in-house. We crossed transgenic mice in which tamoxifen-dependent recombinase Cre^ERT2^ is expressed in an activity-dependent manner from the loci of the immediate early gene Arc (Arc-CreERT2 or B6.Cg-Tg(Arc-cre/ERT2)MRhn/CdnyJ, Jackson Laboratory strain #022357) with transgenic mice expressing a floxed-stop ChR2-EYFP cassette (Ai32 or Ai32(RCL-ChR2(H134R)/EYFP; B6.129S-*Gt(ROSA)26Sor^tm32(CAG-COP4^*^∗^*^H134R/EYFP)Hze^*/J, Jackson Laboratory strain #024109). In offspring possessing both transgenes (Arc-ChR2 mice), neurons in which Arc is induced shortly after tamoxifen injection permanently express ChR2. The specific target immediate early gene (IEG) *Arc* was selected because its expression in HPC is reported to be particularly sensitive to situational context (Guzowski et al. 2001; Chiaruttini et al. 2025). In our broad HPC inhibition experiments we additionally used VGAT-ChR2 mice (B6.Cg-Tg(Slc32a1-COP4*H134R/EYFP)8Gfng/J, Jackson Laboratory strain #014548). In all cases, mouse genotypes were determined by PCR analysis of ear punch DNA samples.

N=16 Arc-ChR2 mice and N=4 cre-negative littermate controls were used for the primary optogenetic perturbation-only experiments, and N=4 Arc-ChR2 mice were used for combined perturbation-electrophysiology experiments. Additionally, we used N=4 cre-negative controls for the fixed-context-cue behavioral control experiment, and N=4 VGAT-ChR2 for the broad optogenetic inhibition control experiment.

One Arc-ChR2 mouse was excluded following data collection due to overall chance performance during laser-off trials during stimulation sessions. No additional mice completed optogenetic perturbation sessions and were subsequently excluded. However, N=16 additional mice were pretrained on the ProAnti task but excluded because i) they failed to reach performance criteria for neuronal tagging (N=11; see *Behavioral Shaping* below), ii) died shortly following tamoxifen treatment (N=3) or NPX implantation (N=1), or iii) had a broken optical fiber cannula (N=1).

Mice were single-housed, or co-housed with same-sex littermates, and maintained on a 12-hour light / dark cycle. All surgical procedures and behavioral training occurred in the dark cycle. All procedures were conducted in accordance with National Institute of Health guidelines and were reviewed and approved by the Institutional Animal Care and Use Committee at Princeton University.

### Surgical procedures

Mice underwent surgical procedures as previously described (Pinto et al. 2018; Cho et al. 2025). In all cases, mice were allowed to recover from each individual surgery for at least 5 days before starting water restriction for behavioral training. Surgery was performed on mice under aseptic conditions and body temperature was maintained with a heating pad (Harvard Apparatus). Mice were anaesthetized with isoflurane (3% for induction, 1–1.5% for maintenance) and given a preoperative intraperitoneal dose of meloxicam for analgesia (10 mg kg^−1^) in warm saline, and a postoperative dose subcutaneously 24 h later. After asepsis, the skull was exposed, and the periosteum was removed.

A custom lightweight titanium headplate was attached to the skull with adhesive cement (C&B Metabond, Parkell). For the optogenetic perturbation experiments without simultaneous electrophysiological recordings, craniotomies over the DG (mediolateral (ML) ±2 mm from the midline; anteroposterior (AP) -1.8 mm posterior from bregma) were made using a pneumatic drill. Chemically-sharpened (Hanks et al. 2015) optical fibers (fibercables.com) were then implanted bilaterally targeting dorsal DG (dorsoventral (DV): 2.0 mm from brain surface at an angle of 10° toward the midline). Another layer of cement was added to attach the fiber cannula to the skull and the headplate.

Procedures were similar for the N=4 mice intended to undergo simultaneous chronic electrophysiological recordings. One mouse received an optic fiber implant as previously described. The other three mice were instead bilaterally implanted with either a single low-profile dual-fiber cannula (Doric TFC_200/250-0.66_3mm_LPB3.0(P)_C60), or two mono-fiber cannulae (Doric MFC_200/240-0.22_2.5mm_ZF1.25_C60) targeting bilateral dorsal DG (ML, ±1.5 from midline; AP, -1.9 from bregma; DV: 1.6mm from brain surface, 0° angle). For all four mice, a gold-coated ground pin (Newark 82K7797) was also implanted through a craniotomy at the cerebellum, approximately 6.0 mm posterior and 2.0 mm lateral from bregma on the left side and affixed with cement. Two mice were implanted with the neuropixel probe prior to training, and two were implanted after pre-training but before tagging.

In the two mice implanted after pre-training, during the same surgery as fiber implantation, fiducial marks were made on the skull over mPFC (ML, ±0.4 mm from the midline; AP, 1.8 mm anterior from bregma). Transparent Metabond (C&B Metabond, Parkell) was used over the fiducial marker to allow visualization of the markers during subsequent craniotomy and probe placement. After training, mice were taken off water deprivation for approximately 1 week until they reached a minimum weight of 25g. In a second surgery, mice were again anesthetized with isoflurane and the Metabond and skull was shaved down with a pneumatic drill. A craniotomy was made over the mPFC at the fiducial marker. A Neuropixel 2.0 probe (phase 2A, four-shank, Imec, N=3; phase 2B, four-shank, Imec, N=1), enclosed in a custom 3D printed casing, was slowly (5 µm/sec) descended through mPFC (DV: 3.0mm from brain surface). The probe was oriented such that the four shanks formed a sagittal plane, aimed at sampling the anterior-posterior extent of the mPFC. Prior to insertion, probe tips were coated with CM-DiI (Invitrogen C7000) dissolved in isopropyl alcohol to 1 µg/µl. Following insertion, Dowsil (3-4680, Dow Corning) was used to cover the craniotomy as a dural sealant. The probe’s ground and reference pads—shorted together prior to surgery—were soldered to the gold pin previously implanted through the skull above cerebellum. The probe assembly was attached to the skull and headplate using Absolute Dentin (Parkell S305), which was then coated with opaque Metabond. The entire NPX assembly was wrapped in Coban for protection until recording. Procedures were the same in two mice implanted prior to all training, except that the entire mPFC craniotomy and NPX probe implant procedure was done during the same surgery as optic fiber implantation.

### Neuropixel implant construction

A custom 3D printable holder was modified from a previous holder used in rats (Luo et al. 2020). All parts were 3D printed in-house using Formlabs SLA printers (Form 3, Form 4) in Grey V4 and Grey V5 Formlabs resin, except for the probe itself and the screws. Printable STL files, as well as assembly instructions including a list of useful tools and links to vendors can be found at https://github.com/Brody-Lab/mouse_npx_holders.

### VR behavioral training

General methods for VR behavioral training are as previously described (Pinto et al. 2018). In brief, mice were head-fixed so that they could run comfortably on an 8-inch (20 cm) Styrofoam® ball suspended by compressed air. Ball movements were measured with optical flow sensors (ADNS3080) via an Arduino Due, and the VR environment was projected onto a coated toroidal Styrofoam® screen (approximately 270° horizontal and 80° vertical visual field) using a DLP projector (Optoma HD141X). The setup was enclosed within a custom-designed cabinet built from optical rails (Thorlabs) and lined with sound-attenuating foam sheeting (McMaster-Carr).

The choice of material and size of the spherical treadmill was designed to minimize the amount of effort to turn the floating ball, such that the moment of inertia of a mouse pushing back the ball (2.78 × 10^−4^ kg m^2^) is comparable to the moment of inertia of a mouse pushing itself (2.68 × 10^−4^ kg m^2^). An optical flow sensor (ADNS-3080 APM2.6), located beneath the ball and connected to an Arduino Due, ran custom code to transform real-world ball rotations into VR movements within the Matlab-based ViRMEn software engine (http://pni.princeton.edu/pni-software-tools/virmen). The ball and sensor of each VR rig were calibrated such that ball displacements (*dX* and *dY*, where X and Y are parallel to the anterior-posterior and medial-lateral axes of the mouse, respectively) produced translational displacements proportional to ball circumference in the VR environment of equal distance in corresponding X and Y axes.

Rewards were delivered by a solenoid valve (NResearch), controlled by a NI USB-6501 (National Instruments). Reward was delivered by sending a TTL pulse to solenoid valves, which were generated according to behavioral events on the ViRMEn computer. Each TTL pulse resulted in the release of a drop of reward (∼4–8ul of 10% sweetened condensed milk in water v/v; or 10% sucrose in water) to a lick tube. Sound was played over speakers located in front of the animal below the toroidal screen.

### ProAnti behavioral task

VR environments were designed within the ViRMEn software engine. The final stage of ProAnti task training consisted of two different 210 cm (100 cm rule cue zone, 100 cm contextual memory delay zone, 10 cm arms) VR T-maze contexts. The two contexts were distinguished based on salient visual textures and colors in the rule cue zone, as well as sky color. The sky color faded to black as the animal ran down the rule cue zone. Different auditory pure tones (Pro: 7 kHz vs. Anti: 12 kHz) played for 50 ms in each context at the start of the trial. After mice exited the rule cue zone, tall, high-contrast visual cues (turn “guide”, 2 towers, 30 cm tall and 2 cm wide each) appeared in one arm of the VR corridor, either to the left or right. Whether the guide appeared on the left or right on a given trial was pseudorandom, with the constraint that motor side biases (left vs. right) were corrected using a debiasing algorithm, as described elsewhere (Pinto et al. 2018). After the memory delay, mice were presented with a liquid reward for turning into the arm with the guide in the Pro context and without the guide in the Anti context. Rewarded trials were followed by a 3.5-s inter-trial interval, and error trials were followed by a 11-s inter-trial interval. During the iti, the visual display froze for the first second after which the display was then blacked out. The median trial length was 10.97±4.0 seconds (median ± 1 m.a.d.). The context switched across trials once mice achieved 70% correct over the previous 20 trials and maintained a motor choice bias (difference in percent correct between right- and left-rewarded trials) below 15% over the previous 20 trials.

*Fixed-context-cue behavioral task variant.* The fixed-context-cue behavior task was identical to the ProAnti task, except that the visual and auditory contextual cues were fixed for the entire duration of the experimental session. Contextual cues were randomly selected prior to the start of each session. The decision rule switched according to the same criterion as for the ProAnti task.

### Behavioral shaping

Following post-operative recovery, mice were extensively handled while gradually restricting water intake to an allotted volume of 1–2 mL per day, delivered either during behavioral sessions or supplemented after sessions. Throughout water-restriction mice were closely monitored to ensure no signs of dehydration were present and that body mass remained at least 80% of the pre-restriction value. Mice were then trained in 1-hr sessions daily on the ProAnti task, according to the shaping procedure described below and in **Fig. S1**. Mice underwent 7 shaping stages (T1-T8), where T8 is the final maze explained in the previous section.

The first 2 shaping stages (T1-T2) consisted of Pro context trials, with progressively increasing lengths. The starting maze was Pro for all mice and not counterbalanced, because pilot studies found that mice took longer to learn the ProAnti task, after context switching was introduced, if they started training in the Anti context. In addition to the turn guide, in mazes T1-T4, there was an additional salient visual turn cue (the “moon beacon”), which was a grey sphere (40 cm diameter) located directly over and behind each trial’s correct choice arm (i.e., the moon beacon was on the same side as the guide in Pro and the opposite side of guide in Anti). In T1-T2, both the moon beacon and the guide were present for the entire trial duration. Mice transitioned from T1 to T2 once they completed 10 trials within a session, with a trial rate of at least 2 trials/min. Mice transitioned from T2 to T3 once they completed at least 40 trials within a session, with a trial rate of at least 2 trials/min and maintained a motor side bias below 20% over the previous 40 trials.

As mice transitioned from T3-T5, the rule cue zone got shorter, and the memory delay zone got longer. The final length of the rule cue and memory delay zones (described in the previous section) was achieved in T5. Context switching across blocks was introduced in T3. In all mazes with context switching (T3-T8), mice switched between contexts once they achieved 70% correct over the previous 20 trials. Additionally, mice had to maintain a motor side bias below 10% in T3-T4, and below 15% in T5-T8.

In T3-T4, the first trial of each context block had the moon beacon present at the start of the trial. We reasoned that as mice learned to go to the side with the moon beacon, they incidentally learned the context-specific decision rules (Pro = toward the guide; Anti = away from the guide) as well. However, on subsequent trials, the moon beacon appeared with increasing delay relative to the mouse’s VR starting position as the mouse ran down the maze stem, according to an ascending 1-up/1-down adaptive staircase procedure yoked to the mouse’s previous within-block performance. In other words, as mice performed well, the moon would appear with increasing delay as they run down the maze stem, and vice versa if they are performing poorly. The moon appeared on trial t (for all t>1) when mice ran through a position along the maze stem (in cm), according to:

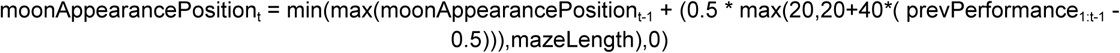

where prevPerformance_1:t-1_ is the percentage correct over all previous trials within the block, and mazeLength was the entire length of the VR maze. For t=1, moonAppearancePosition_t_=0. If moonAppearancePosition_t_ was equal to the maze length, the moon did not appear during that trial. This gradual offset of the moon beacon relative to trial start was designed to cause mice to attend to the more stable indicator of correct choice (the turn guide) rather than the transiently available moon.

Mice transitioned from T3 to T4 once they completed 2 context switches with at least 20 trials each within a session, with a trial rate of at least 2 trials/min and a motor side bias below 10% over the previous 20 trials. Mice transitioned from T4 to T5 if they completed 2 context switches within a session, with an average performance greater than or equal to 80% correct in the final block and a motor side bias below 10% over the previous 20 trials.

T5 was the same as T4, except there was no moon beacon on the first trial. Rather, on t=1, moonAppearancePosition_1_=mazeLength, so that if mice made an error, the moon would appear at decreasing delays relative to the mouse’s VR starting position. Each context block in T5 began with a warmup in T3. Note that during these warmup trials, and all subsequent mazes (T6-T7) with warmup trials, the length of the rule cue and memory delay zones were inherited from the mouse’s current maze (and not the same as T3). The warmup maze had a minimum of 10 trials, and once mice achieved 60% correct performance in the warmup maze, mice started T5. Mice transitioned from T5 to T6 once they completed 2 context switches within a session with an average performance greater than or equal to 70% correct in the final block and a motor side bias below 15% over the previous 20 trials.

In T6, each context block began with a warmup maze in T3. This warmup maze had a minimum of 5 trials, and once mice achieved at least 3/5 correct performance in the warmup maze, they transitioned to T6, which was the same as T5 except there was no moon. If mice performed below 55% correct over the previous 20 trials, they received a 5-trial long ‘easy’ block in T3 with the moon. Mice transitioned from T6 to T7 if they made 2 context switches within a session, with an average performance greater than or equal to 70% correct, a trial rate of at least 2 trials/min and a motor side bias below 15% over the previous 20 trials. Mice who completed T6 met the criteria for a neuronal tagging session (see *Neuronal tagging* section below). T7 was the same as T6 except that the guide only appeared after mice left the rule cue zone. Mice completed a 5-trial long warmup maze in T6 in T7 (55% correct to transition). Mice transitioned from T7 to T8 if they made 2 context switches within a session, with an average performance greater than or equal to 70% correct, a trial rate of at least 2 trials/min and a motor side bias below 15% over the previous 20 trials. T8 was the final stage of training described in the last section, which was the same as T7 except that there was no warmup maze.

### Neuronal tagging

Recombination was induced in mice via an i.p. injection of 4-hydroxytamoxifen (4-OHT; 200 μl/mouse at 10 mg/ml). 4-OHT powder was first mixed with 100% ethanol (15 mg/375 μl) and vortexed vigorously. The solution was then poured into a 50°C chamber with corn oil, vortexing for approximately 1h until fully dissolved, to create a stock solution that was stored at −20° C until required. On experimental days, aliquots of the stock solution were heated to 37° C prior to injection.

Approximately 5.5 hours following 4-OHT treatment, mice were trained for a single session in a single context (either Pro or Anti, counterbalanced across mice). This tagging session occurred after mice transitioned out of maze T6 during behavioral shaping, and before T7. Like T6, each tagging session began with a 5-trial T3 warm up maze, followed by T6 but with no context switching. Unlike T5 during behavioral shaping, there were no demotions during the tagging session to T3 based on behavior. Following the tagging session, mice were left undisturbed for 72 hours to prevent off-target neuronal tagging. During this isolation period, mice only interacted with an experimenter to receive their daily water allotment and get weighed in the colony. After this 72-hour isolation period, mice continued behavioral shaping on maze T6 for at least one session to ensure their behavior fully recovered post-tagging prior to transitioning to T7.

The tagging procedure for the Dark-tag control mice was like that of the ProAnti mice. However, approximately 5.5 hours following 4-OHT treatment, Dark-tag mice were placed on the treadmill in the dark in the VR rig for 1 hour and received rewards at random intervals with an average reward frequency matching that with which they received rewards during T5 behavioral shaping.

### Optogenetics during VR behavior

Following behavioral shaping and the tagging session, mice underwent optogenetic testing. Bilateral optical fibers were coupled to a 473-nm DPSS laser (LaserGlow, 200 mW) via fiberoptic patch cables (Doric, MFP_062/125/900-0.22_1m_FCM-MF2.5 or SBP(2)_200/220/LWMJ-0.22_1m_FCM-2xMF2.5). For optogenetic perturbation experiments without simultaneous electrophysiological recordings, mice received laser illumination (∼2 mW, measured from the output of a pre-implanted fiber; 15 ms pulses) on 30% of trials that was restricted to either the rule cue zone only (0–100 cm; 10% of trials), the memory delay region only (100–200 cm; 10% of trials), or the entire maze stem (0-200 cm; 10% of trials). Laser power was titrated during an independent pilot experiment (data not shown). In all experiments with Arc-ChR2 mice or Cre- controls, optogenetic stimulation occurred at 20 Hz. While it is possible to tag and artificially express context-specific behaviors via photo-stimulation of CA1—and thus it is possible that behavioral biases in the present experiments may in part reflect off-target effects—this has been previously observed at 4 Hz, not at 20 Hz (Ryan et al. 2015). In experiments with VGAT-ChR2 mice, stimulation occurred at 40 Hz. For each mouse, we performed 3-5 individual optogenetics sessions (mode: 5). Five sessions were selected as the maximum number because an independent pilot experiment revealed behavioral biases induced by stimulation peak during the first week of optogenetics testing, with more variable results subsequently, potentially due to ongoing learning.

For optogenetic perturbation experiments with simultaneous electrophysiological recordings, mice also received laser illumination on 30% of trials. Unlike the opto-only mice described above, however, perturbation trials in these mice were restricted to the rule cue zone (15% of trials) or the memory delay (15% of trials) of the T-maze (i.e., these mice did not receive entire-maze-stem-stimulation trials). During the first experimental session for each simultaneous opto-ephys mouse, data from which were excluded from further analysis, we confirmed that the laser did not cause salient photostimulation artifacts on mPFC voltage traces during optical stimulation; no visible stimulation artifacts were observed at powers <6 mW.

To confirm that HPC laser stimulation did not directly stimulate mPFC neurons, we performed a control experiment in N=3 opto-ephys mice (**Fig. S13**). This control experiment was conducted following all other data collection for each mouse. In this control experiment, mice performed the ProAnti task during which brief single pulses of laser light (10ms duration) were delivered on every trial at random locations of the T-maze stem. These data were then used to estimate the latency of response of mPFC neurons following laser stimulation, compared to simulated data in which neurons were directly activated by stimulation. One simultaneous opto-ephys mouse did not perform this control experiment, and was excluded from all latency analyses; however, complementary analysis of this mouse’s main-text optogenetic experiments yielded similar results as those shown in **Fig. S13**.

The laser was controlled by TTL pulses generated using a National Instruments DAQ card on a computer running the ViRMEn-based virtual environment. The TTL signals were fed to a TTL Pulse Generator (Doric; OPTG_4 or OPTG_8) to create and gate the desired light pulse trains. The latency between the Matlab command to send the TTL trigger on the ViRMEn computer and laser light output was measured to always be <6ms.

### Electrophysiological recordings during VR behavior

Neuropixel data was acquired at 30 kHz. Raw voltage traces were filtered, amplified, multiplexed, and digitized on-probe and recorded using SpikeGLX (https://billkarsh.github.io/SpikeGLX). The VR computer emitted a binary pattern of seven independent TTL streams (NI USB-6501) that encoded the ViRMEn frame number, and these pulses were recorded in SpikeGLX using an auxiliary National Instruments data acquisition card (NI PXIe-6341 with NI BNC-2110) to synchronize behavioral and neurophysiological data. This card and the Imec card were placed in the same PXIe-1071 chassis. VR data were synchronized to spiking data by computing the current VR frame number from the pattern of TTL pulses recorded simultaneously to the NPX data.

Upon reaching behavioral criteria for optogenetic experiments, each neuropixel implanted mouse’s electrophysiological signal was inspected. To target the ACC, we selected a channel map of active recording sites on a mouse-by-mouse basis. We first identified the location of the brain surface along the probe shanks, based on the lack of local field potential and observable spiking during the task. We then constructed a channel map consisting of a horizontal stripe across all four shanks, with 96 active sites per shank, aligned dorsally approximately 0.5 millimeters ventral from the brain surface. In all cases, this channel map was additionally consistent with the expected distance from the probe tips, given our total insertion depth of 3 millimeters. Although a fixed channel map was then used for all further recording sessions for each mouse, the signal was qualitatively re-inspected at the start of each recording day to confirm it remained stable.

### Behavioral data analysis

Trials with response latencies (i.e., time to make a choice) greater than three standard deviations above the median response time were excluded from all analysis (2.8% of all trials), regardless of experimental group, laser or trial condition. These trials typically involved backtracking along the stem of the maze.

#### ProAnti task behavior

Behavioral performance was first quantified as the percentage of trials in which mice turned toward (TT) the turn guide, corresponding to the use of the *pro* decision rule (**Fig. 1E**). For Pro-and Anti-tagged mice, as well as Cre- controls, this measure was then converted to reflect the percentage of trials in which mice used the tag-session-consistent decision rule: for Anti-tag mice, this was computed as 1 – TT (i.e., % of trials mice turned away from the guide); for Pro-tag-session mice, it was simply TT. Thus, higher values indicated greater use of the tagged rule.

To assess the rule bias induced by optical stimulation (e.g., **Fig. S4**), we compared the expressed decision rule during laser-on trials (TT_on_) to a surrogate distribution of matched laser-off trials (TT_off_). This surrogate distribution was generated via bootstrap resampling (K=1000) separately for each mouse, matching laser-off trials to laser-on trials by number of trials, time since context switch (binned into tertiles: early, middle, late), and choice. Rule bias was defined as TT_on_ minus the mean of the TT_off_ distribution for Pro-tagged mice, and as TT_off_ minus TT_on_ for Anti-tagged mice, ensuring that positive bias values consistently reflected increased use of the tagged rule during stimulation. To compute the decision rule bias separately for each choice, the procedure was repeated separately for each type of choice, now only matching the number of trials and time since context switching. Significance of the observed bias for each session was assessed by comparing TT_on_ to the bootstrap distribution of TT_off_ values, using percentile rank. Results were qualitatively similar including previous outcome variables such as previous reward or previous choice in the generation of the surrogate TT_off_ distribution.

Motor bias was computed as the absolute difference in performance between turn-guide-left and turn-guide-right trials. We assessed the motor bias induced by optical stimulation in a similar manner as the decision rule bias, comparing motor bias during laser-on trials to a surrogate distribution of matched laser-off trials. This surrogate distribution was generated via bootstrap resampling (K=1000), matching laser-off trials to laser-on trials by number of trials, time since context switch (binned into tertiles: early, middle, late) and previous choice.

Velocity and heading angle were analyzed across the maze stem to examine effects of HPC engram reactivation on decision-rule-irrelevant factors (**Fig. S5**). For each trial, position and linear velocity were binned along the maze stem (40 bins), and average velocity or heading angle per bin was computed. Laser effects were quantified as the difference between laser-on and -off trials for each mouse, context (Tag vs. Untag), and trial outcome (Correct vs. Error). Statistical significance was assessed using a paired bootstrap procedure: mice were resampled with replacement, and for each resampled mouse, trials were resampled with replacement (K=10,000). The mean difference across resampled trials was computed for each iteration, producing bootstrap distributions of paired differences from which two-sided p-values were calculated. P-values were FDR-corrected for multiple comparisons across maze bins and trial outcomes per context.

#### Fixed-context-cue behavior

We assessed differences between task variants (ProAnti vs. Fixed-context-cue) along three different behavioral measures—trials to criterion, number of context switches per session, and task performance—using within-subject permutation tests that accounted for repeated measures across sessions within mice. For each measure, session-level values were first averaged within mice to obtain condition means, and the observed mean difference between conditions was computed across mice. To assess significance, condition labels were shuffled across sessions within each mouse (K=1,000), preserving the number of sessions and blocks per session, and the mean difference between conditions was recalculated for each permutation to generate a null distribution. P-values were defined as the proportion of permuted mean differences with absolute values greater than or equal to the observed difference.

### Electrophysiological data analysis

#### Preprocessing

Neuropixel data was preprocessed with CatGT (https://github.com/billkarsh/CatGT) to apply global common average referencing (-gblcar) and to isolate the action potential frequency band (-apfilter=butter,12,300,9000). Spike sorting was then performed offline using Kilosort4. This pipeline was implemented in datajoint (https://www.datajoint.com/) and ran automatically over newly generated files: New files produced by SpikeGLX were automatically copied to an in-house cluster using Globus. On this cluster, equipped with GPUs, a set of Slurm scripts automatically ran CatGT, Kilosort, and custom synchronization code. When sorting and synchronization was complete, spike times and cluster identities were extracted from the output of KiloSort and Phy using code from the Spikes repository (https://github.com/cortex-lab/spikes). We used Phy to identify clusters with obvious irregularities such refractory-period violations or high noise contamination to classify clusters as single units (“good”). Finally, clusters with mean firing rates below 0.2 Hz across all ProAnti trials were discarded. All unit isolation and curation were performed blind to the session’s corresponding behavioral data.

Within each session, average spike counts over time for each unit were computed in non-overlapping 10 ms windows. Spike counts were smoothed over time with a 100ms causal filter. To facilitate averaging across trials of different lengths, each trial was interpolated to be 250 timepoints long, such that the rule cue and memory delay were each 100 timepoints, the time in the arms was 50 timepoints. The ITI was excluded from further analysis.

#### Single unit analyses

To provide an overall description of single-unit encoding of context, turn guide side, and choice during laser-off trials (**Fig. S8B**), we computed each unit’s average firing rate during each correct laser-off trial, separately for the rule cue and memory delay epochs. For each unit, we then fit a generalized linear model (GLM) to the epoch-averaged trialwise firing rates with context, turn guide side, and choice as predictors, including interaction terms, and tested whether each factor explained significant firing rate variance (p<0.05, uncorrected), separately for the rule cue and memory delay epochs. These statistical thresholds were used for basic characterization and as an inclusive criterion for downstream analyses, rather than for population-level inference; all primary statistical conclusions rely on permutation-based and cluster-corrected analyses described below.

For coding direction analyses described below (e.g., **Fig. 3**), we restricted analysis to the population to neurons that encoded any of these task variables (or any combination) during any considered task epoch, as only these neurons can reliably contribute to changes in task representations during HPC reactivation. The same population of neurons was used for all coding direction analyses (e.g., choice-only modulated neurons were included in context coding direction analyses and vice versa). Results were qualitatively identical when using the full population.

We also examined how mPFC single-unit activity was modulated by task context, choice, and laser perturbation over time (e.g., **Fig. 4**). All comparisons of time-resolved firing rate traces were performed on matched trial sets to control for behavioral covariates. Trial matching was performed independently for each neuron and repeated across K independently resampled matched trial sets per neuron (K=10 for laser-off analyses and K=50 for laser-on analyses). For context and choice selectivity, trials were matched on choice and context, respectively. For laser-on versus -off comparisons within a given context, trials were matched on choice. Additionally, in laser-on analyses, laser-off trials were further resampled to match the pre-laser mean baseline firing rate of laser-on trials, ensuring that baseline differences did not drive observed laser effects.

Condition differences were quantified using time-resolved Cohen’s d. For each matched resample, Cohen’s d was computed as the difference in mean firing rates between conditions divided by the pooled standard deviation at each time point. These effect-size traces were then averaged across matched resamples to yield a single time-resolved effect size per neuron. Statistical significance was assessed using a nonparametric permutation approach. Within each matched resample, trial labels were randomly permuted between conditions (B=100 permutations), and null Cohen’s d values were recomputed at each time point. Two-tailed permutation p-values were calculated at each time point as the proportion of permuted values exceeding the observed effect in absolute value. This procedure yields a resample-conditioned null distribution that respects the trial-matching structure.

For context and choice, neurons were considered selective only if permutation p-values exceeded a threshold of p<0.01 at any time point. Neurons were classified by preference (e.g., Tagged vs. Untag context preferring; left vs. right choice preferring) based on the sign of their maximal absolute Cohen’s d during laser-off trials. Neurons without significant selectivity were considered non-preferring. Neurons were sorted by the time of peak effect size and categorized as rule cue or memory+arm field cells depending on whether the peak occurred during or after the rule cue epoch.

To quantify laser effects within each context and field location, Cohen’s d time courses were computed separately for laser-on and laser-off conditions using the same matched-trial procedure. Laser effects were then assessed by comparing these resample-averaged effect-size traces. Population-level uncertainty was estimated using hierarchical bootstrapping across mice and sessions. For each bootstrap iteration (K=5,000), mice were resampled with replacement, sessions were resampled within mice, and neurons were resampled within sessions. For each bootstrap sample, the median Cohen’s d across neurons was computed at each time point. Population traces are reported as the median across neurons, with variability estimated from the bootstrap distribution.

To test whether population response to the laser over time differed from laser-off baseline while correcting for multiple comparisons, we used a nonparametric cluster-based permutation test (Maris and Oostenveld 2007). For each time point, the test statistic was the median across neurons. The null distribution was generated using a sign-flipping permutation procedure in which the sign of each neuron’s response was randomly inverted and the median time course recomputed. P-values were calculated as the proportion of permuted medians whose absolute value exceeded the observed median. Time points exceeding a two-sided cluster-forming threshold (α=0.05) were grouped into contiguous temporal clusters. Cluster-level statistics were defined as the sum of the absolute median values within each cluster. Significance was determined by comparing observed cluster statistics to a permutation distribution of the maximum cluster statistic, thereby controlling the FWE rate. Clusters with p<0.01 were considered significant.

#### Coding direction analysis

We quantified low-dimensional population structure by computing coding directions (CDs) that maximally discriminated activity patterns associated with context, turn guide side, and choice. For a population of n simultaneously recorded neurons, the mean firing-rate vectors for Tagged vs. Untagged contexts (x̄_tag,t_, x̄_untag,t_), left vs. right guide side (x̄_guide left,t_, x̄_guide right,t_), or left vs. right choice (x̄_left,t_, x̄_right,t_) were computed at each time point t from laser-off trials. For context, the trial-by-trial difference w_t_^context^ = x̄ – x̄ was stable across laser-off trials but varied across task epochs (**Fig. S9**). Thus, the context axis was defined at each time point, smoothed within each epoch using a 100-bin moving mean and normalized at each t. For guide and choice, we followed prior work (Inagaki et al. 2019) in defining fixed coding directions over task periods during which the corresponding signals were stable. For guide, the difference w_t_^guide^ = x̄ – x̄_guide right,t_ was not stable prior to the appearance of the turn guide; therefore, the guide axis was defined as the average of w_t_^guide^ over the memory delay, divided by its Euclidean norm. For choice, the difference w_t_^choice^ = x̄ – x̄ was stable across laser-off trials during the end of the memory delay but not prior to the appearance of the turn guide. Therefore, the choice axis was defined as the average of w_t_^choice^ over the second half of the memory delay, divided by its Euclidean norm.

To avoid imbalances in the trial distributions when estimating the CDs, we performed a re-balancing procedure repeated 100 times. For each subsampling iteration, we constructed two independent matched sets of laser-off trials (Set 1 and Set 2), each balanced across the four context–choice combinations (tagged/untag × left/right). For each CD type, we computed two coding directions (CD1 and CD2), one from each matched set. Critically, all projections were performed using a held-out CD: trials belonging to Set 1 were projected using CD2, and trials belonging to Set 2 were projected using CD1. This ensured that CD estimation and projection were statistically independent on every iteration.

Single-trial population trajectories were obtained by projecting firing-rate vectors onto the relevant CD using the inner product. For each session, trialwise trajectories were centered at each time point by subtracting the mean projection across all correct laser-off trials, and then normalized by the maximum absolute projection within that session to enable pooling across sessions. Standard errors were estimated via hierarchical bootstrapping: sessions were sampled with replacement, and within each sampled session, trials were resampled with replacement (K=1,000).

We quantified neural projection differences across context (Tagged vs. Untag), guide (left side vs. right side), choice (left vs. right), or laser status (on vs. off) using a hierarchical bootstrap analysis. Analyses were performed on predefined temporal windows (10 timebins wide each), with each trial’s projection values averaged across time points within each window. For each window, we performed a hierarchical bootstrap (K=2000): resampling subjects with replacement, selecting one session per resampled subject, and resampling trials with replacement within each condition. Separation between conditions was quantified using the area under the ROC curve (AUC) within each bootstrap draw. For each iteration, a null distribution was generated by randomly permuting condition labels (e.g., laser on/off), and observed AUC values were compared to their null counterparts in a paired manner to obtain two-sided bootstrap p-values. P-values were corrected for multiple windows tested. Similar results were obtained using Cohen’s *d* and Jensen-Shannon divergence to quantify distributional differences.

To determine how much population activity variability was captured by the context axis, we computed the fraction of across-neuron variance explained by the context axis on a trial-by-trial basis, separately for laser-off and -on trials. For each cross-validation shuffle, the context axis was evaluated only on held-out trials. For each trial, the population activity *X* (neurons x time) was first centered across neurons at each timepoint. At each timepoint t, the component of the activity vector X_t_ within the context axis was computed as 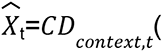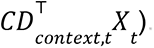. Residual activity for computing variance was then 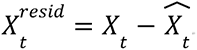 Within each analysis window (rule cue zone or memory delay), we computed the total activity variance as the sum across neurons of each neuron’s variance across time within the window (SST), and the corresponding residual variance after removing the context axis projection (SSE). The variance explained by the context axis on that trial was given by 1-SSE/SST. This metric can take negative values when the context axis derived from held-out data fails to generalize to a given trial, analogous to cross-validated R^2^ in regression. For comparison, a null distribution was obtained by randomly rotating the context axis relative to the neural population activity with orthogonal rotation matrices prior to projection onto the context axis, and recomputing variance explained (K=50 per shuffle).

#### Decoding of task variables from population activity

To quantify how mPFC population activity predicted task variables on a trial-by-trial and timepoint-by-timepoint basis, we applied linear discriminant analysis (LDA) to decode decision rule (pro vs. anti), choice (left vs. right), context (Tagged vs. Untag), and turn guide from neural activity at each time point.

LDA models were trained using only laser-off trials. To ensure balanced sampling, training trials were grouped into condition bins defined by combinations of task variables. For rule decoding, bins were defined by context, choice, rule, and trial outcome (correct, error). For decoding of other variables, bins were defined by choice, context, and turn guide side. The smallest bin size determined the number of trials sampled per condition. For each of 10 iterations, an equal number of trials per bin was randomly selected to construct a balanced training set.

At each time point, population activity across neurons was z-scored using the mean and standard deviation of the training data. LDA models were fit using a pseudo-linear discriminant with L2 regularization (MATLAB *fitcdiscr*). For rule decoding, the shrinkage parameter (gamma) was optimized independently at each time point using Bayesian optimization. For decoding of the other task variables, to reduce computational load, gamma was optimized at a middle time point within each epoch (rule cue, memory delay, arm) and then held fixed across all time points within that epoch.

At each time point, LDA models were trained using the balanced training set and applied to all remaining trials (including laser-on trials). For each trial and time point, the decoder returned posterior class probabilities. A decision variable (DV) was defined as the difference between class probabilities, and predicted labels were assigned based on its sign. Trials used for training were excluded from evaluation within each iteration. Decoding outputs were averaged across iterations. Population decoding accuracy was computed at each time point as the fraction of correctly classified trials. In addition, for each trial, we quantified the time of peak decoding magnitude (peak |DV|).

Statistical comparisons were performed using hierarchical bootstrap resampling, in which mice were sampled with replacement, followed by resampling of sessions and trials within each mouse. Bootstrap distributions were used to test decoding accuracy against chance (50%) and to compare conditions (e.g., laser-on vs. laser-off, correct vs. error trials), separately for each task variable.

#### Hidden Markov Modeling of context dynamics

We used a two-state hidden Markov model (HMM) to infer discrete latent "context states" from single-trial population trajectories projected along the context axis, pooling projections across mice and sessions (**Fig. S14**).

The HMM was first fit to laser-off correct trials to estimate emission parameters (state-specific means μ and variances σ) using a linear emission model that allowed for asymmetric or sloped means over time. Transition probabilities (A) were then fit to memory-delay reactivation trials with emissions fixed, using the expectation–maximization (EM) algorithm with weak Dirichlet priors (α = 0.05). To prevent baseline offsets in laser-on trials from producing artifactual transitions, context axis projections during the first half of the rule-cue zone were recentered to laser-off medians, and this pre-laser period was excluded from transition analyses. Because trials were analyzed independently rather than as a continuous sequence, this adjustment ensured that each trial started in the appropriate contextual state without constraining the HMM’s initial state, allowing transition probabilities to be inferred naturally within each trial.

For each trial, the posterior probability of being in each state (γ) was computed and used to classify trials according to their pre-laser state. To identify the timing of laser-evoked transitions, we computed a first-state-shift index using a sliding window analysis. For each trial, the model identified the first time bin within the memory delay in which the posterior crossed from the pre-laser dominant state to the opposite state (e.g., state tag → untag for initially tagged-state trials). Trials without a detected crossing were classified as "no shift." To exclude ambiguous cases, we removed trials where the mean posterior following the detected shift (+5 bins) was inconsistent with a genuine state transition (i.e., mean γ > 0.5 for tag → untag transitions, or < 0.5 for untag → tag transitions). The proportion of trials exhibiting a detectable transition was computed as the fraction of identified shifts among all trials starting in a given pre-laser state.

Laser-evoked state-transition latencies were converted to seconds using the same interpolated time-from-trial-start alignment applied to firing rates. These transition times were used to examine guide and choice coding conditioned on the contextual state transition. Specifically, we aligned timepoint-by-timepoint LDA guide or choice decoding accuracy, as described previously, for each reactivation trial to its inferred contextual state transition time. Decoding accuracy on each trial was baseline-corrected by subtracting mean decoding in the pre-shift window (bins -400ms to 0ms).

To construct a null distribution that preserved global temporal trends in decoding within each trial (e.g., ramping) while breaking the link between neural data and transition time, we repeated this alignment using permuted shift times (K=1000). On each permutation, each shift trial was assigned a transition time borrowed from a different shift trial, with the constraint that the donor trial’s transition time differed from the target trial’s transition time by at least 400ms. Each trial’s own decoding data was then extracted at the permuted transition time and baseline-corrected using the same pre-shift window at the permuted time. Permuted traces were averaged across trials to yield one null observation per permutation, producing a null distribution of 1000 mean peri-transition decoding profiles. To summarize the post-shift divergence between real and null, we computed a z-score over the 1000–1600ms post-shift window as the mean difference (real − null) divided by the standard deviation of the null distribution over the same window.

#### Spectral analysis of 20 Hz entrainment

All spectral analyses (**Fig. S16**) were performed on binned spike counts in non-overlapping 10ms bins, within a window of 1 second preceding, and 3 seconds following, memory delay zone laser onsets. Similar results were obtained with 1ms bins. For initially assessing the presence of 20 Hz power, we performed multitapering analyses on each cell and trial independently using the *spectrum* library in Python (Cokelaer et al., 2017). The multitapering analyses were averaged over the first four prolate spheroidal sequences with a time half bandwidth of 2.5. Trials in which mice backtracked more than 5% of the total maze length were excluded. Individual trial-cell pairs in which the cell failed to fire more than 2 spikes were excluded (∼5% trial-cell pairs). We excluded any trial in which fewer than 5 cells passed this threshold (∼0% trials). For comparison, we also obtained a similar window of time aligned to the onset of the memory delay zone in all laser-off trials, and computed an identical multitapering analysis. For each cell we then averaged the power spectrum above 4Hz across all valid laser-on or laser-off trials. For **Fig. S16A** the average across cells was normalized to [0,1] to ease comparison.

We searched for peaks in each cell’s average power spectrum using the *find_peaks* function from the *scipy* library in Python (Virtanen et al. 2020). Peaks were defined as the highest local maxima within a window of 30Hz that were at least 1 standard deviation above its lowest contour line, generally resulting in 0-3 peaks per cell. Example cells were pulled from the set of all cells that possessed their most prominent peak in their trial-averaged PSD at 20 Hz, where prominence refers to the distance to a peak’s lowest contour line.

To understand the timing of 20 Hz entrainment across each trial, we used the continuous wavelet transform (CWT) function from the *pywavelet* library in Python (Lee et al. 2019). We used a complex Morlet wavelet centered at 100Hz with a bandwidth of ∼67Hz. Similar results were obtained using other wavelet functions. Because we were interested in power in the 20 Hz band, we considered the wavelet transform at scales corresponding to log-spaced frequencies from 4Hz to 40Hz. For all analyses we computed the squared absolute magnitude of the complex result, and we discarded a padding of 500ms at the edges of our time window to mitigate warping due to cone-of-influence effects.

To ensure accurate estimates of power, we discarded low firing rate cell-trial pairs similarly to the multitapering analyses; however, given the time-resolved nature of the CWT, we also discarded any cells that failed to fire *on average* two spikes within the entire memory delay zone across all considered trials (∼40% cells). We then computed the CWT for each valid cell and trial independently, before averaging spectrums across all cells. Drawing on methods from the EEG field (Pfurtscheller and Lopes da Silva 1999), we averaged power within each frequency band across the 500ms baseline preceding laser onset (or in the case of matched trials, preceding the memory zone onset) to convert energy to the percent change from baseline. To quantify 20 Hz entrainment, we translated these values to be nonnegative and then took the ratio of mean percent change within the [18.5, 21.5] Hz band to mean percent change across all other frequencies within the [4, 40] Hz range.

Statistical comparisons between laser-on and laser-off trials followed the methodology of (Durka et al. 2004), with the exception that we generated our null distribution from an equal number of laser-off trials restricted to match the distribution of left and right choices from the laser-on trials. Statistical tests accounted for multiple comparisons and time-frequency bin multiplicity by controlling the false discovery rate to 1% and performing permutation tests with 100,000 repetitions. Heatmaps of matched laser-off trials are computed from our null distribution averaged across all repetitions. Error bars for plotting the 20 Hz ratio were computed as the standard deviation of a randomly resampled laser-on distribution across all repetitions.

#### Spatiotemporal sequence analyses

We quantified the similarity of spatiotemporal activity sequences observed in mPFC using two complementary approaches. First, to assess sequence similarity between laser-off and laser-on trials in the untagged context while removing firing-rate amplitude differences, we compared population activity vectors elicited during laser-off and laser-on trials. For each neuron, we computed the average activity across trials, and then normalized each neuron’s activity by its maximum across time, separately for laser-off and laser-on trials. Neurons without detectable spiking in either condition were excluded. At each time point, sequence similarity was then computed as the Pearson correlation across neurons between laser-off and laser-on population vectors. A noise ceiling was estimated using trial-matched subset correlations within laser-off trials: for each of 100 iterations, laser-off trials were randomly partitioned into two non-overlapping subsets each matched in number to the laser-on trials, each subset was independently normalized by its own per-neuron maximum across time, and sequence similarity was calculated between subsets. A rotational control was generated by randomly shifting neuron identities within the laser-off activity matrix and recomputing similarity to laser-on activity (K=100 iterations).

Second, we implemented a Bayesian decoder (**Fig. S17**). For each session, trials were divided into laser-off trials from the Untag context, laser-on trials from the Untag context, and laser-off trials from the Tagged context. Trial counts were matched across conditions by subsampling to the minimum available number. Decoding was repeated across multiple (K=10) random, non-overlapping train/test splits within each session, and results were averaged to obtain a single estimate per session.

Decoding performance was evaluated by training and testing across distinct trial sets. For each split, tuning curves were computed from Untag context laser-off training trials as the mean firing rate of each neuron at each time bin and smoothed with a causal half-Gaussian kernel (σ=2 bins). Decoding was performed using a Poisson-like Bayesian model, with decoded spatiotemporal position estimated as the position bin maximizing the log-likelihood of observed activity. Within-context, cross-laser, and cross-context decoding was assessed by applying the Untag laser-off trained decoder to held-out Untag laser-off trials, Untag laser-on trials, and Tagged laser-off trials, respectively. Performance was quantified as the median absolute error (MAE) between decoded and true position at each time point, along with signed decoding error (decoded minus true position). Statistical comparisons were performed across sessions using paired nonparametric tests.

### Histology

Mice were deeply anesthetized (i.p. 200 mg/kg ketamine and 20 mg/kg Xylazine) and transcardially perfused with 1x phosphate-buffered saline (PBS), followed by fixation with 4% paraformaldehyde (PFA). Whole brains were post-fixed in 4% PFA for 1–2 days, and then placed in a 30% (v/v) sucrose solution (1 x PBS, 0.1% Sodium Azide). ChR2-EYFP expression, as well as fiber optic and NPX probe placement, were assessed using light sheet imaging. PFA-fixed brains were washed several times, cleared and stained according to the iDISCO+ protocol (Renier et al. 2014, 2016); and as previously described in (Zimmerman et al. 2025), and used to acquire high-resolution whole-brain light sheet imaging volumes.

Specifically, brain samples were serially dehydrated in increasing concentrations of methanol (Carolina Biological Supply, 874195; 20%, 40%, 60%, 80% and 100% in doubly distilled water (ddH2O); 45 min–1 h each), bleached in 5% hydrogen peroxide (Sigma, H1009) in methanol overnight and then serially rehydrated in decreasing concentrations of methanol (100%, 80%, 60%, 40% and 20% in ddH2O; 45 min–1 h each).

Brain samples were washed in 0.2% Triton X-100 (Sigma, T8787) in PBS, followed by 20% DMSO (Fisher Scientific, D128), 0.3 M glycine (Sigma, 410225) and 0.2% Triton X-100 in PBS at 37 °C for 2 days. Brains were then washed in 10% DMSO and 6% NDS (EMD Millipore S30) and 0.2% Triton X-100 in PBS at 37 °C for 2–3 days to block nonspecific antibody binding. Brains were then washed twice for 1 h at 37 °C in 0.2% Tween-20 (Sigma P9416) and 10 mg ml–1 heparin in PBS (PTwH solution) followed by incubation with primary antibody solution (chicken anti-GFP, 1:500; Aves labs, 1020) in 5% DMSO, 3% NDS and PTwH at 37 °C for 7 days. Brains were then washed in PTwH 6 times for increasing durations (10 min, 15 min, 30 min, 1 h, 2 h and overnight) followed by incubation with secondary antibody solution (Alexa Fluor 647 donkey anti-chicken, 1:500; Invitrogen, A78952) in 3% NDS and PTwH at 37 °C for 7 days. Brains were then washed in PTwH 6 times for increasing durations again (10 min, 15 min, 30 min, 1 h, 2 h, overnight).

Brain samples were serially dehydrated in increasing concentrations of methanol (20%, 40%, 60%, 80% and 100% in ddH2O; 45 min–1 h each), then incubated in a 2:1 solution of dichloromethane (Sigma, 270997) and methanol for 3 h then washed twice for 15 min in 100% dichloromethane. Before imaging, brains were stored in the refractive-index-matching solution dibenzyl ether (Sigma, 108014).

The 488 channel was used to align the imaged volumes to the Allen atlas (https://github.com/cortex-lab/allenCCF) and confirm optic fiber placement; CM-DiI-related fluorescence at 561 was used to infer electrode tracks. Track tracing (**Fig. S3C-D**) was performed using BrainGlobe tools (Tyson et al. 2022) and visualized with Brainrender (Claudi et al. 2021). Expression was confirmed in the 639 channel.

Due to previously reported issues with arcTRAP YFP staining in whole cleared brains (Pavlova et al. 2018; Franceschini et al. 2025), we focused on groupwise comparisons of gross fluorescence (ProAnti vs. Cre-; Dark-tag vs. Cre-; Dark-tag vs. ProAnti). Fluorescence intensities were normalized within-subject by dividing the registered fluorescence volume by the mean fluorescence measured inside the ventricles (defined based on the Allen atlas) and an out-of-brain region. Group comparisons were restricted to the hippocampal region of interest (ROI), defined as the union of CA1-3 and dentate gyrus regions from the Allen atlas. Within this ROI, we defined spherical sampling regions (radius = 5 voxels) and retained only spheres whose centers overlapped the ROI by at least 25%. Sphere locations were identical across subjects. For each sphere and subject, we computed the mean fluorescence across all voxels in the sphere. Group differences at each sphere were assessed using a Mann-Whitney U test (one-sided for comparisons with Cre- and two-sided for comparison between Dark-tag and ProAnti-tag), with FDR-correction applied across spheres.

### General statistics and Reproducibility

Optogenetics experiments, but not analysis, were conducted blind to experimental conditions. All experiments were conducted over multiple cohorts. Individual cohorts consisted of a random selection of test groups (i.e., Pro-tag, Anti-tag, Cre-negative, Dark-tag mice). We did not account for cohort effects in our statistical analyses, but did not observe obvious cohort-dependent effects. No statistical method was used to predetermine group sample sizes; rather, animal and session numbers were chosen based on independent pilot studies as previously described (data from which are not included here).

---

**Table S1.** Statistical tests in Figures 2, 3, and S4.

| Panel | Comparison | Test | Statistic or 95% CI | df | p-value |
| --- | --- | --- | --- | --- | --- |
| 2B | Pro-tag mice, Tag: laser-on vs. off | Paired t-test | t = 0.86 | 5 | 0.79 |
| 2B | Pro-tag mice, Untag: laser-on vs. off | Paired t-test | t = 24.77 | 5 | 3.0×10<sup>-6</sup> |
| 2B | Pro-tag mice, Untag: laser-on vs. chance (50%) | One-sample t-test | t = 5.36 | 5 | 0.002 |
| 2B | Anti-tag mice, Tag: laser-on vs. off | Paired t-test | t = 0.51 | 5 | 0.69 |
| 2B | Anti-tag mice, Untag: Laser-on vs. off | Paired t-test | t = 4.39 | 5 | 0.004 |
| 2B | Anti-tag mice, Untag: laser-on vs. chance (50%) | One-sample t-test | t = 2.49 | 5 | 0.028 |
| 2B | Pro-tag, Context x Laser | rm-ANOVA | F=93.34 | 1,5 | 0.0002 |
| 2B | Anti-tag, Context x Laser | rm-ANOVA | F=11.35 | 1,5 | 0.02 |
| 2C | PA mice, Untag: laser-on vs. off | Paired t-test | t = 9.499 | 11 | 1.23×10<sup>-6</sup> |
| 2C | PA mice, Untag: laser-on vs. chance (50%) | One-sample t-test | t = 5.088 | 11 | 0.0004 |
| 2C | PA mice, Tag: laser-on vs. off | Paired t-test | t = 0.917 | 11 | 0.379 |
| 2C | PA mice, Context × Laser | rm-ANOVA | F = 50.585 | 1,11 | 2.0×10<sup>-5</sup> |
| 2C | PA mice, laser-on (Untag) vs. laser-off (Tag) | Paired t-test | t = 1.71 | 11 | 0.116 |
| 2D | Cre-, Tag: laser-on vs. off | Paired t-test | t = 1.58 | 3 | 0.213 |
| 2D | Cre-, Untag: laser-on vs. off | Paired t-test | t = 1.98 | 3 | 0.143 |
| 2D | Cre-, Context × Laser | rm-ANOVA | F = 3.74 | 1,3 | 0.15 |
| 2E | Absolute motor-choice bias (all groups) | One-sample t-test | ts < 0.467 | 11 or 3 | > 0.65 |
| 2E | All group comparisons | Unpaired t-tests | ts < 0.811 | 14 | > 0.43 |
| 2F | PA mice, left choices | One-sample t-test | t = 2.477 | 11 | 0.015 |
| 2F | PA mice, right choices | One-sample t-test | t = 10.246 | 11 | 2.90×10<sup>-7</sup> |
| 2F | Cre- controls, left or right choices | One-sample t-tests | ts < -0.736 | 3 | > 0.74 |
| 2F | PA vs. Cre-, left | Welch's t-test | t = 2.572 | 13.57 | 0.023 |
| 2F | PA vs. Cre-, right | Welch's t-test | t = 9.051 | 9.48 | 5.71×10<sup>-6</sup> |
| 2F | Tagged context, PA or Cre-, left or right | One-sample t-tests | ts < 1.26 | 11 or 3 | > 0.14 |
| 2G | Dark-tag, Pro: laser-on vs. chance (50%) | One-sample t-test | t = 0.226 | 3 | 0.84 |
| 2G | Dark-tag, Anti: laser-on vs. chance (50%) | One-sample t-test | t = 1.82 | 3 | 0.17 |
| 2G | Dark-tag, Pro: laser-on vs. off | Paired t-test | t = 3.21 | 3 | 0.025 |
| 2G | Dark-tag, Anti: laser-on vs. off | Paired t-test | t = 3.59 | 3 | 0.019 |
| 2G | Dark-tag, Context × Laser | rm-ANOVA | F = 43.98 | 1, 3 | 0.007 |
| 3M | Tagged - laser-off & laser-on, decoding > chance (2nd half of memory delay) | Permutation test | Off CI = [0.703–0.927]<br>On CI = [0.673–0.920] | – | both <0.001 |
| 3M | Untagged - laser-off & laser-on, decoding > chance (2nd half of memory delay) | Permutation test | Off CI = [0.609–0.950] | – | both <0.001 |
|  |  |  | On CI =<br>[0.544–0.903] |  |  |
| 3M | Tagged, laser-on vs. off, (2nd half of memory delay) | Permutation test | CI =<br>[-0.127–0.073] | – | 0.578 |
| 3M | Untagged, laser-on vs. off, (2nd half of memory delay) | Permutation test | CI =<br>[-0.156–0.038] | – | 0.256 |
| 3N | Corr on > off (middle half of memory delay, bins 125-175) | Hierarchical bootstrap | CI = [-0.0087<br>1.2774] | – | 0.026 |
| 3N | Err on > off (middle half of memory delay, bins 125-175) | Hierarchical bootstrap | CI = [0.0246<br>1.1124] | – | 0.015 |
| 3N | Corr on > off (end of memory delay, and arms, bins 176-250) | Hierarchical bootstrap | CI = [-0.0783<br>0.6806] | – | 0.058 |
| 3N | Err on > off (end of memory delay, and arms, bins 176-250) | Hierarchical bootstrap | CI = [0.0477<br>1.0892] | – | 0.016 |
| S4B (Left) | Anti-tag bias in Pro context | One-sample t-test | t = 4.883 | 5 | 0.002 |
| S4B (Left) | Pro-tag bias in Pro context | One-sample t-test | t = -0.954 | 5 | 0.808 |
| S4B (Left) | Cre- bias in Pro context | One-sample t-test | t = -1.79 | 3 | 0.914 |
| S4B (Left) | Anti- vs. Pro-tag | Unpaired t-test | t = 4.694 | 10 | 0.0008 |
| S4B (Left) | Anti-tag vs. Cre- | Unpaired t-test | t = 4.285 | 8 | 0.0027 |
| S4B (Left) | Pro-tag vs. Cre- | Unpaired t-test | t = 0.068 | 8 | 0.948 |
| S4B (Right) | Pro-tag bias in Anti context | One-sample t-test | t = 20.62 | 5 | $2.48 \times 10^{-6}$ |
| S4B (Right) | Anti-tag bias in Anti context | One-sample t-test | t = -0.892 | 5 | 0.794 |
| S4B (Right) | Cre- bias in Anti context | One-sample t-test | t = -1.863 | 3 | 0.920 |
| S4B (Right) | Anti- vs. Pro-tag | Unpaired t-test | t = 5.392 | 10 | 0.0003 |
| S4B (Right) | Anti-tag vs. Cre- | Unpaired t-test | t = 0.025 | 8 | 0.981 |
| S4B (Right) | Pro-tag vs. Cre- | Unpaired t-test | t = 11.511 | 8 | $2.94 \times 10^{-6}$ |

**Figure S1.**
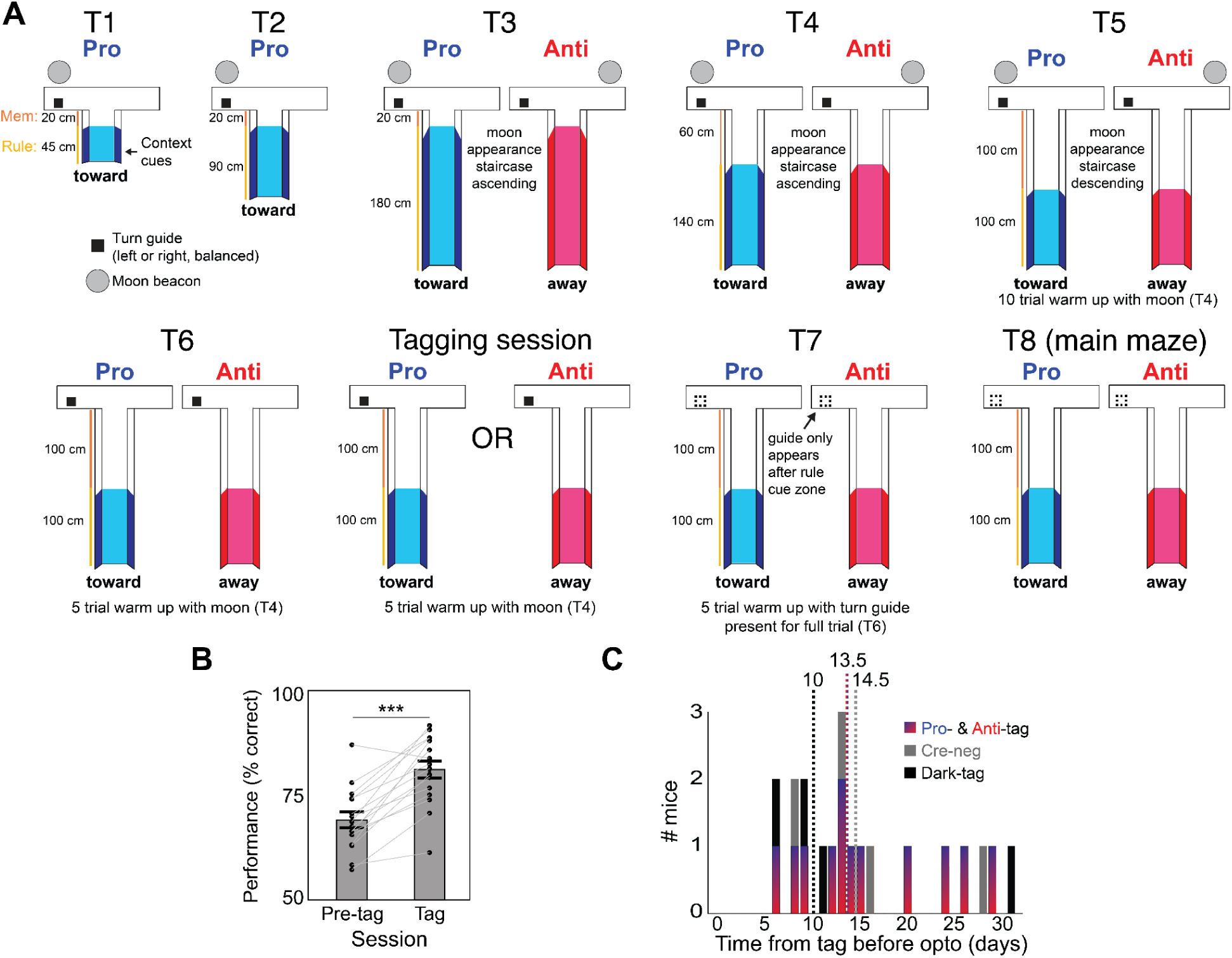
ProAnti task shaping and tagging procedure. **(A)** Schematic of the 7 different shaping mazes (T1 – T7), optional tagging session following T6, and the final ProAnti maze (T8) (*Materials & Methods: Behavioral shaping*). Moon staircase refers to the moon beacon appearing with increasing (or decreasing) delay relative to the mouse’s VR starting position, according to an adaptive staircase procedure yoked to the mouse’s performance within a given context block. When mice made correct choices, the moon would appear with increasing delay as they ran down the maze stem, or vice versa if they made incorrect choices. Ascending moon staircases were such that the moon appeared at the start of the trial on the first trial in a context block, and descending moon staircases were such that the moon beacon did not appear on the first trial of a context block but returned as mice made errors. **(B)** Comparison of performance (% correct) during the tagging session, compared to performance in the same context during pre-tagging session (t-test; Pro- & Anti-tag and Cre- mice; N=16; t_15_=6.73, ***p=6.77x10^-6^). Dots denote individual mice, error bars ±1 s.e.m. **(C)** Number of behavioral training days between tagging session and start of HPC stimulation sessions, by experimental group (dotted lines show group-specific medians).

**Figure S2.**
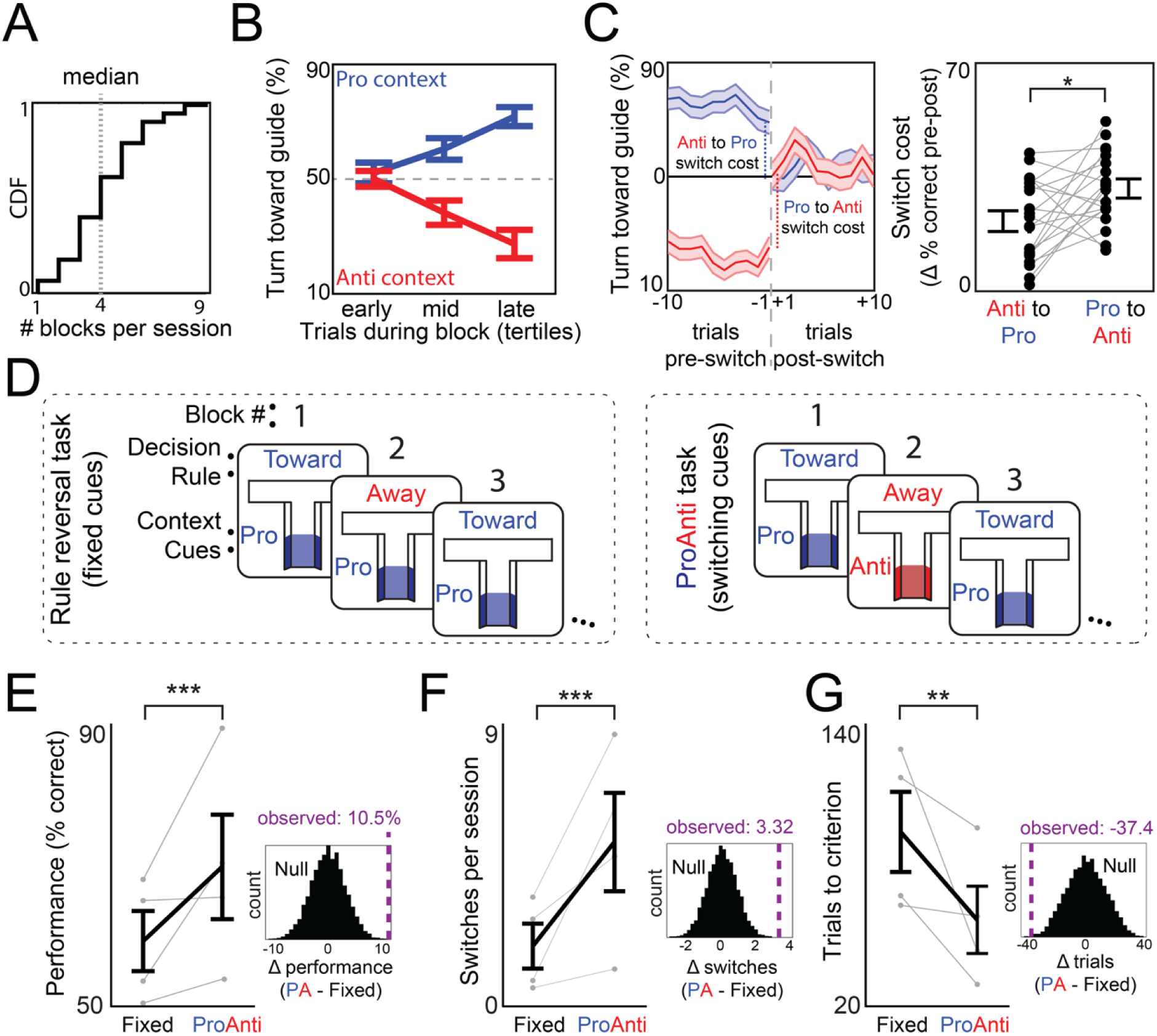
Mice use the contextual cues to switch decision rules, despite context switch cost. **(A)** CDF of completed context blocks per behavioral session (N=90 sessions). **(B)** % trials mice turn toward the turn guide over time in a block, split into tertiles, separately for each context (N=20 mice). Mice showed a performance cost following switches, with weaker performance in the first tertile following context switching than at end of the block. **(C)** *Left*: % trials mice turn toward the turn guide relative to the block switch time, separately for each context. Switching cost was computed as the difference in performance (% correct) between the last ten trials before context switching (pre) and the first ten trials after context switching (post) in the same context. *Right*: There was a smaller switching cost going from Anti to Pro than Pro to Anti (t-test, t_19_=2.79, p=0.012). **(D)** Mice used the contextual cues to switch decision rules across blocks. *Left*: Following ProAnti training, an independent group of mice (N=4) were tested in a rule reversal control task in which the contextual cues were fixed across blocks for the entire session duration, but the decision rule switched according to the same protocol as ProAnti. If mice ignored context cues, this would not affect behavioral performance. *Right*: By contrast, in the ProAnti task, the context cues change in concert with the decision rule (same as Fig. 1B). **(E)** Mice performed worse in the fixed-cue task than in the ProAnti (PA) task (hierarchical permutation test, p<0.00001; see inset). As a result of the impaired performance, **(F)** mice completed far fewer context switches during the fixed-cue task than ProAnti (p<0.00001), and **(G)** took far more trials reach the context-switch criterion during the fixed-cue task than ProAnti (p=0.001). Error bars ±1 SEM. Dots denote mice. *p<0.05, **p<0.01, ***p<0.001

**Figure S3.**
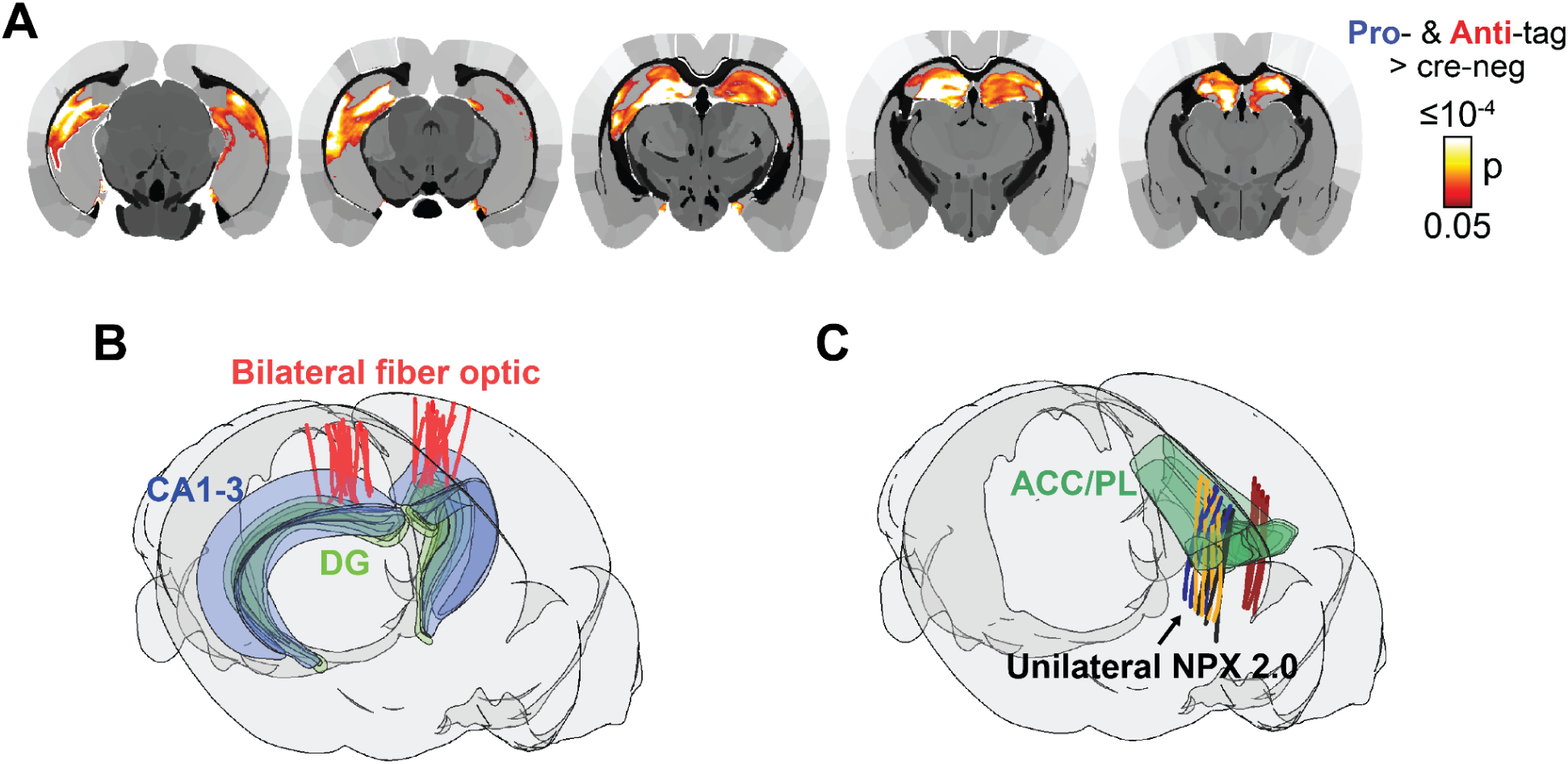
Histology. **(A)** Contrast of ChR2-EYFP expression between Pro- & Anti-tag mice vs. Cre-control mice within a hippocampal region of interest. Pro- & Anti-tag mice exhibited greater expression than Cre- controls (p<0.05, FDR corrected), including in the DG. **(B)** Fiber optic implant locations, targeting DG, in the Pro- & Anti-tag and Cre- mice. (**C**) Four mice were additionally implanted with Neuropixel 2.0 probes with four shanks each in mPFC (Anterior Cingulate Cortex–ACC, Prelimbic Cortex–PL). Track colors denote mouse.

**Figure S4.**
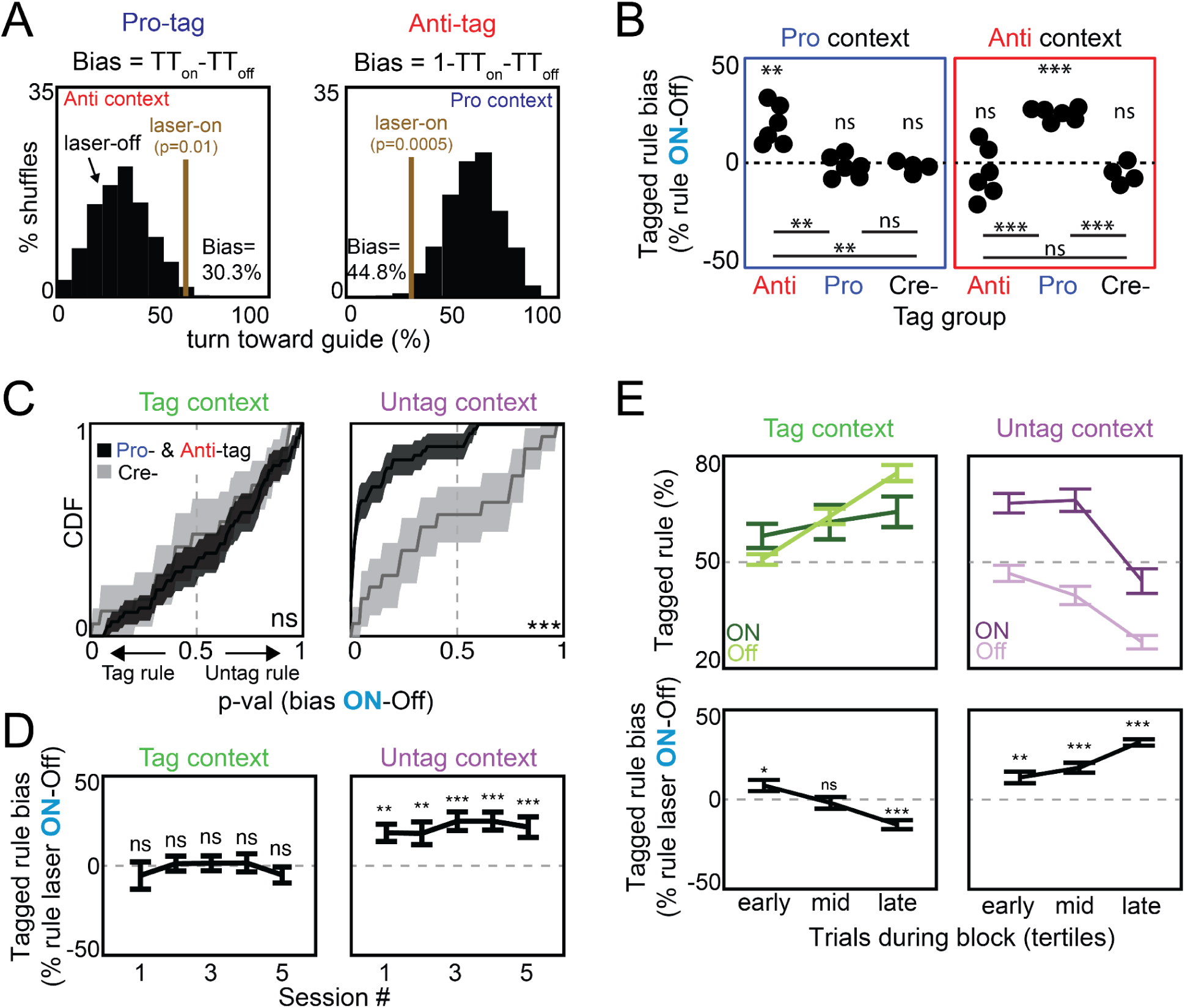
Reliability of context-specific biases induced by HPC engram reactivation. Laser-on trials correspond to memory delay laser epoch in all panels. **(A)** Method for calculating tagged rule bias illustrated for an example session each from one Pro-tag mouse (left) and one Anti-tag mouse (right). Laser-off trials were matched to laser-on trials in terms of previous choice, time in a block since context switching, and number of trials, via bootstrap resampling (laser-off distribution). The tagged rule bias was the difference in mean of the shuffle laser-on distribution and the observed laser-off turn toward (TT) guide percentage in Pro-tag mice, and vice versa in Anti-tag mice. The significance of the bias was computed based on the percentile rank of the laser-on TT versus laser-off TT distribution. **(B)** Tagged rule bias was context and group-specific: Anti-tag mice were biased in the Pro context (p=0.002), Pro-tag mice in the Anti context (p=0.000002), and Cre- showed no bias. Full Statistics in Table S1. **(C)** CDF of tagged-rule bias p-values across all individual sessions in Pro- and Anti-tag (N=12; PA) and Cre- (N=4) mice, separately in the Tagged (left) and Untag (right) contexts. PA exhibited stronger tag-specific biases than Cre- in the untagged (KS test, p=0.0003) but not tagged context (p=0.81). **(D)** PA showed a tagged rule bias during each behavioral session in the Untagged context (one-tailed t-test, all ts>2.79, ps<0.009), but not Tagged (all ts<0.31, ps>0.38), with no dependence on session number in either context. **(E)** *Top*: PA tagged rule (% trials turn in the tagged direction) over time in a block, split into tertiles (independent of laser status) separately for each context (Tagged vs. Untag) and laser status (Off vs. On). *Bottom*: Rule bias over time in block, computed as in (A), but separately for each tertile. In the Untag context, PA showed a tagged rule bias independent of time in a block since context switching (one-tailed t-tests; all t_11_s>3.81, ps<0.003). In the Tagged context, PA showed a tagged-rule bias early in a block after context switching (two-tailed t-test, t_11_=2.51, p=0.03) – when mice perseverated with the untagged rule during laser-off trials – but no bias in the middle of the block (t_11_=0.57, p=0.58). Performance was biased away from the tagged rule at the end of the block (t_11_=5.59, p=0.0002); however, behavior during this end-of-block time window was still tag-like overall, and the absolute magnitude of this decrease in tag-like behavior relative to laser-off trials was smaller than the magnitude of the corresponding tag rule bias in the Untag context within the same time window (|tagged rule bias late| untag vs. tag context, t_11_=5.57, p=0.0002). Error bars ±1 s.e.m. in (C-D), and ±1 bootstrap SD in (B). *p<0.05, **p<0.01, ***p<0.001

**Figure S5.**
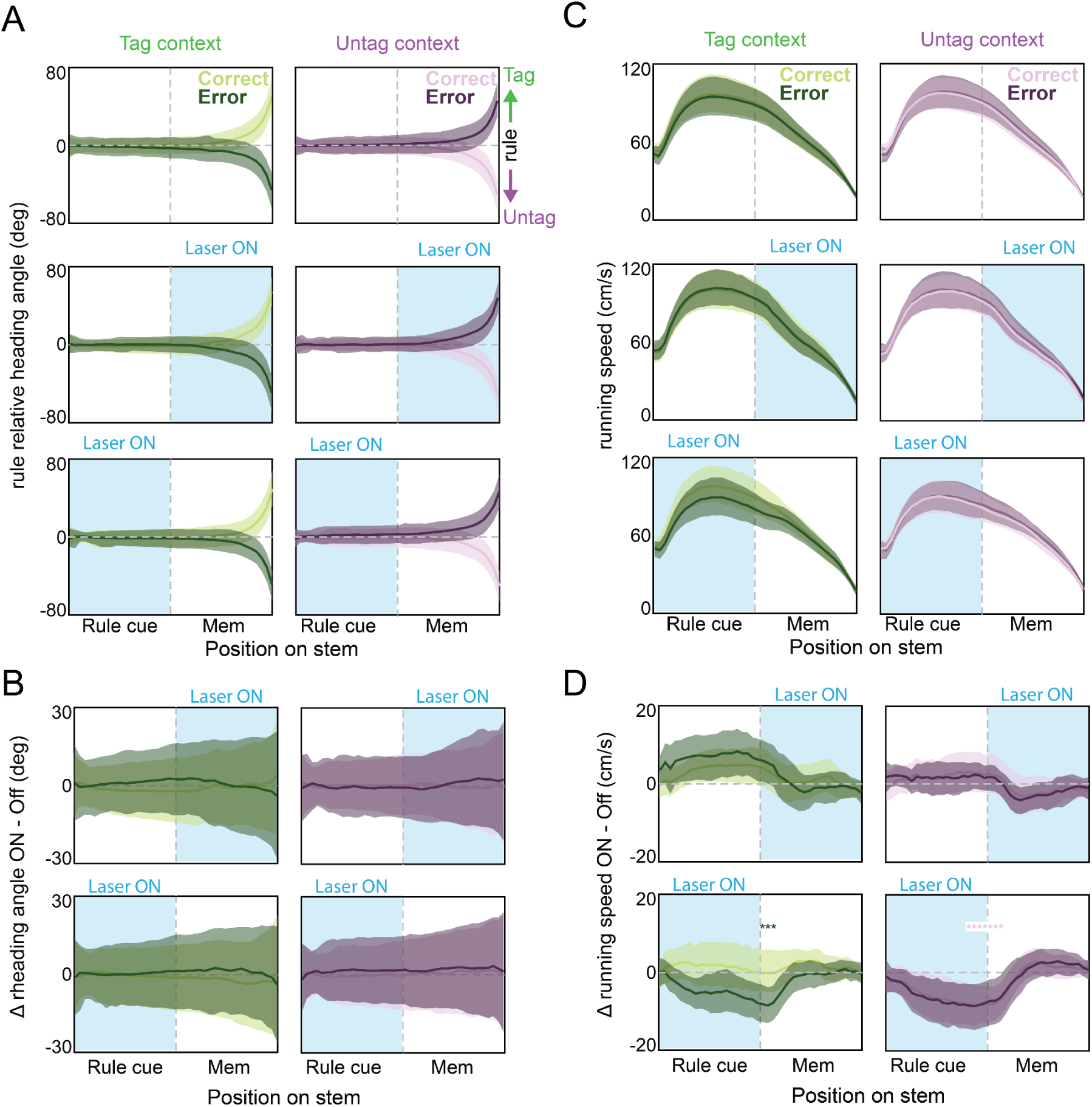
Effect of HPC engram reactivation on decision-rule-irrelevant factors. **(A)** Virtual reality (VR) heading angle across the maze stem in Pro- and Anti-tag mice (N=12), split by context (Tag, left; Untag, right) and trial outcome (Correct, dark; Error, light) for laser-off (top), memory delay reactivation (middle), and rule cue reactivation (bottom) trials. Positive heading angles correspond to directions associated with tag-related rule choices (facing the turn guide in Pro-tag mice, away from it in Anti-tag mice). Negative values correspond to the opposite. **(B)** Bootstrap difference between laser-on and laser-off trials, separated by laser epoch, trial outcome, and context. No significant laser effects were observed. **(C)** Same as (A), but for running velocity across the maze stem. **(D)** Same as (B), but for running velocity. There was no significant effect of memory delay stimulation on running speed. By contrast, mice slowed at the transition between the rule cue zone and memory delay during rule cue reactivation trials when they subsequently expressed the untagged decision rule (i.e., error in Tag context; correct in Untag context). In all plots, lines indicate bootstrap mean ±1 s.e.m. *p<0.05, FDR corrected.

**Figure S6.**
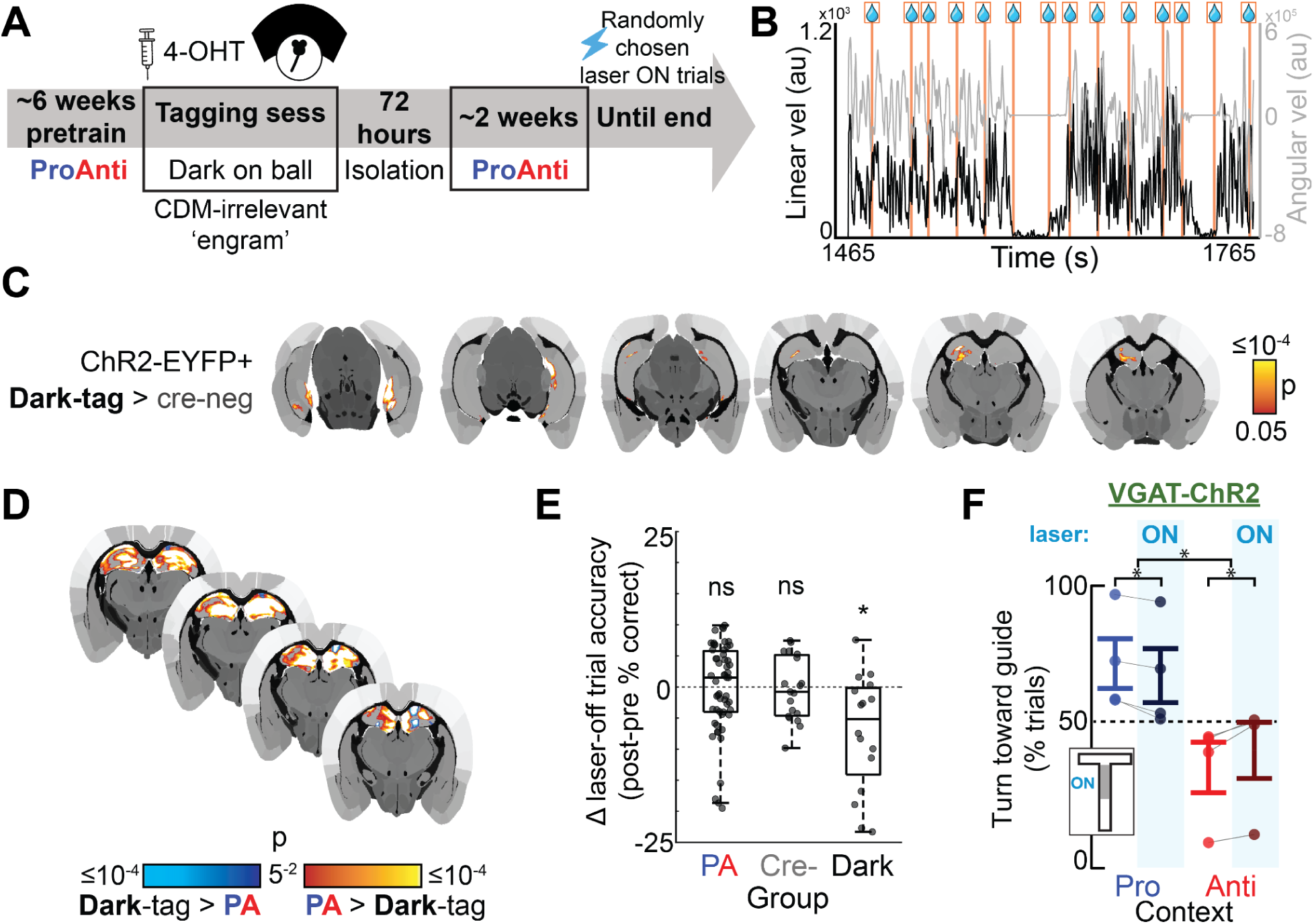
ProAnti-task-irrelevant HPC ensembles. **(A)** Experimental timeline for Dark-tag mice was the same as for Pro- & Anti-tag mice except that following 4-OHT treatment, mice were placed on the ball in the dark and randomly interspersed rewards. **(B)** Linear and angular velocity during a representative 5 minutes of a dark-tag tagging session. Orange vertical lines denote time of reward. Mice tended to run during the entire tagging session, only changing velocity immediately following reward. **(C)** Contrast of ChR2-EYFP expression in Dark-tag vs. Cre- mice. Dark-tag exhibited greater expression than Cre-neg controls (Dark-tag > Cre-neg) in the hippocampal formation (p<0.05, FDR corrected), including in the DG. **(D)** Contrast of ChR2-EYFP expression in Dark-tag vs. Pro- and Anti-tag (PA) mice. PA exhibited greater expression strength than Dark-tag throughout most of the dorsal hippocampus (hot colors). **(E)** Boxplot (median, interquartile range, full range) of difference in performance (% correct) during laser-off trials during sessions in which optogenetic perturbations were performed and average performance during the session prior to perturbations. Dark-tag mice showed a performance impairment (mixed-effects model, p=0.021), but PA and Cre- did not (ps>0.81). Dots denote sessions. **(F)** Bulk inhibition targeting the DG impaired performance on the ProAnti task in VGAT-ChR2 mice (N=4) (Pro, t-test t_4_=4.10, p= 0.03; Anti, t_4_=-4.25, p=0.02; rm-ANOVA, context x laser interaction: F_1,3_=32.47, p=0.01). Dots denote mice, error bars ±1 s.e.m. *p<0.05, **p<0.01, ***p<0.001

**Figure S7.**
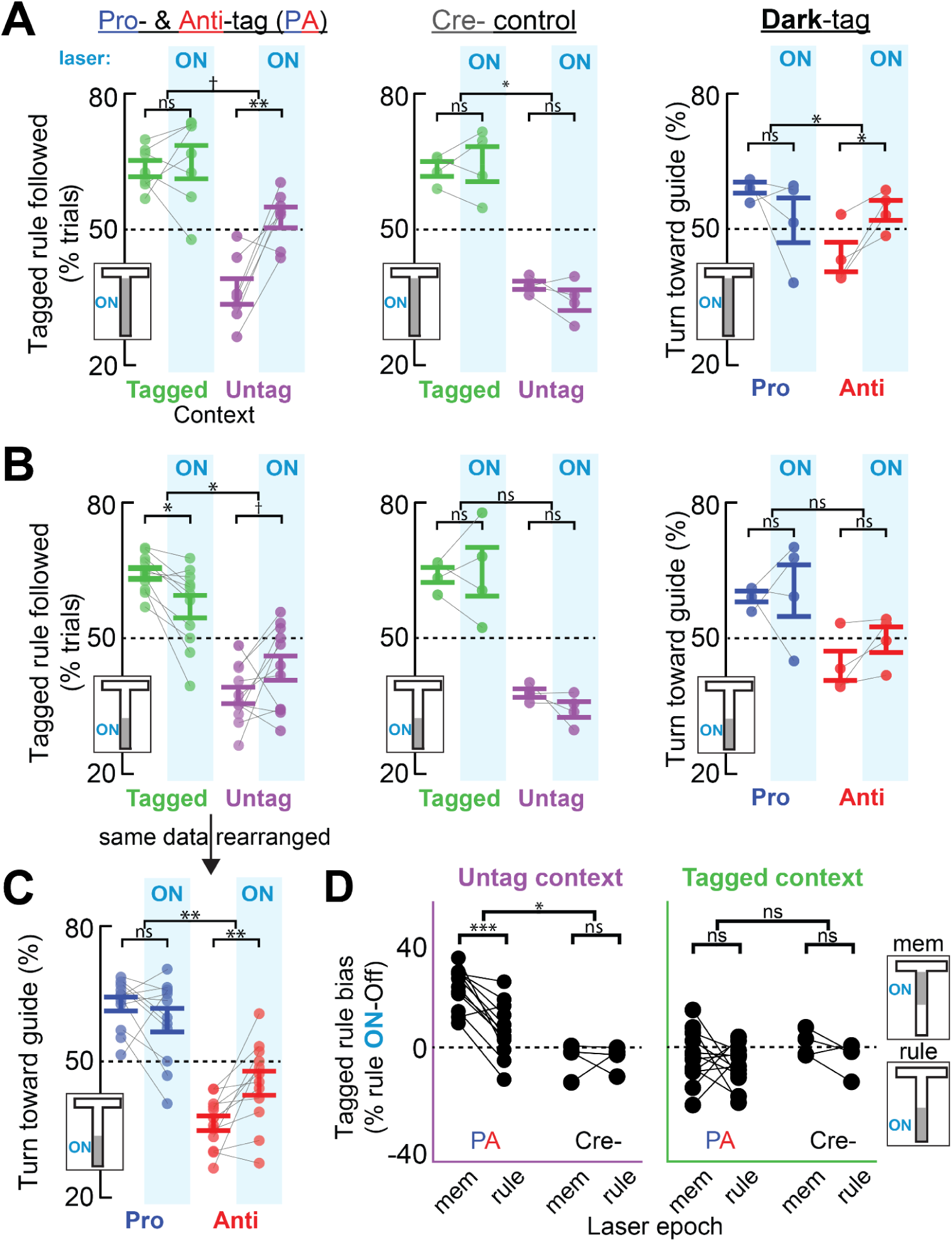
HPC engram reactivation does not evoke tag-specific biases during context recognition. **(A)** Same as Fig. 2B-D, but for trials in which the laser was turned on during the entire VR maze stem (i.e., both the rule cue and memory delay epochs; inset schematic; 10% of trials). *Left*: Results were qualitatively like those in Fig. 2B, in which the laser was turned on only during the memory delay: Pro- & Anti-tag (PA) mice to use the tagged decision more often during laser-on than -off trials in the untagged context (N=7 mice; t-test, t_6_=4.22, p=0.006), but no laser effect was observed in the tagged context (t_6_=0.76, p=0.38) (marginal context x laser interaction: F_1,6_=4.95, p=0.068). Unlike during memory delay reactivation, whole stem reactivation in the Untag context did not elicit as strong an effect as laser-off in the Tagged context: t_6_=3.83, p=0.009. *Middle*: Same as (A, left), but for Cre- mice (N=4). No laser effect in either context (both t_3_<1.6, p>0.21), though the interaction was significant (F_1,3_=11.78, p=0.04). *Right*: Same as Left but for Dark-tag mice (N=4) split by context (Pro, Anti). The laser impaired performance in both contexts, though only significantly in the Anti context (t_3_=3.54, p=0.039), not Pro (t_3_=1.26, p=0.30) (interaction: F_1,3_=19.38, p=0.02). **(B)** Same as (A) but for trials in which the laser was on exclusively in the rule cue zone (inset schematic; 10-15% of trials). *Left*: Rule cue reactivation caused PA (N=12) to perform worse in both Tagged (t_11_=2.92, p=0.014) and Untag (t_11_=2.08, p=0.061) (context x laser interaction: F_1,11_=12.067, p=0.005). *Middle (Cre-) and Right (Dark-tag)*: no laser effect in either context (all t<1.96, p>0.145; interactions: both F_1,3_<1.38, ps>0.33). **(C)** Same data as in (B, left), but resorted by Pro/Anti context rather than Tagged/Untag, independent of tag group. Rule cue zone reactivation impaired performance in Anti context (t_11_=3.60, p=0.004), but not Pro (t_11_=1.42, p=0.19) (interaction: F_1,11_=12.07, p=0.005). **(D)** Comparison of tagged rule bias between memory and rule cue zone laser epochs in PA and Cre-, separately for the Untag and Tagged contexts. Untag context: PA, t_11_=5.58, p=0.0002; Cre-, t_3_=0.04, p=0.97; group x epoch interaction: F_1,14_=8.37, p=0.01. Tagged context: PA, t_11_=0.73, p=0.48; Cre-, t=1.62, p=0.20; interaction: F_1,14_=0.07, p=0.79. Dots denote individual mice. Error bars denote ±1 SEM; ^†^p<0.07, *p<0.05, **p<0.01, ***p<0.001

**Figure S8.**
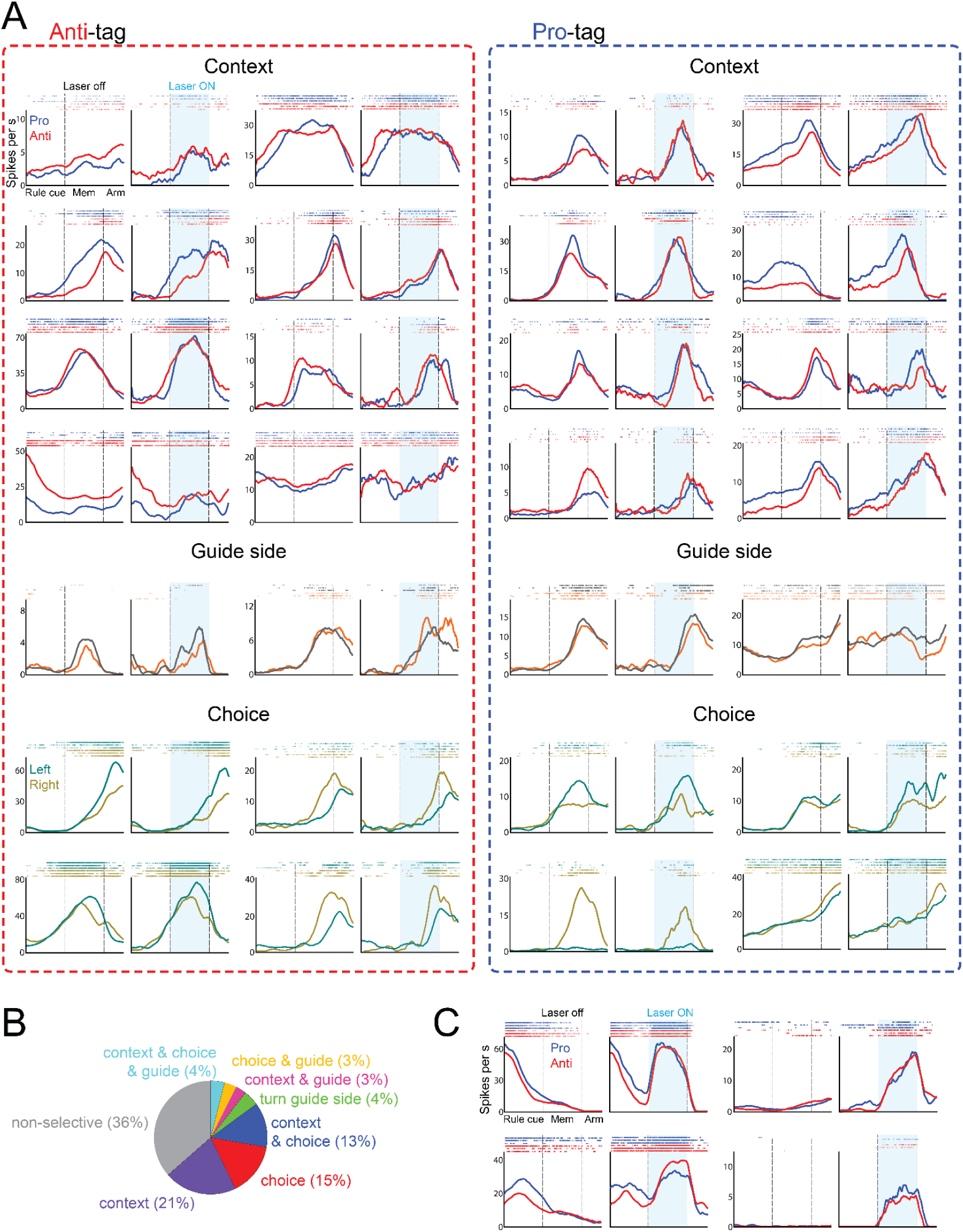
Single units in mPFC represent context, guide, and choice. **(A)** Representative single unit firing fields and associated raster plots (6 random trials) in Anti-tag (left) and Pro-tag (right) mice. Context (top), turn guide side (middle), and choice (bottom) modulated cells, laser-off (left) and -on (right) during the memory delay reactivation. **(B)** Percentage of mPFC cells that encode context, turn guide side, or choice along the VR maze stem (GLM; inclusion threshold: p<0.05). **(C)** Rare examples of single unit firing fields that show strong firing-field-location-independent activity changes during memory delay reactivation compared to laser-off trials.

**Figure S9.**
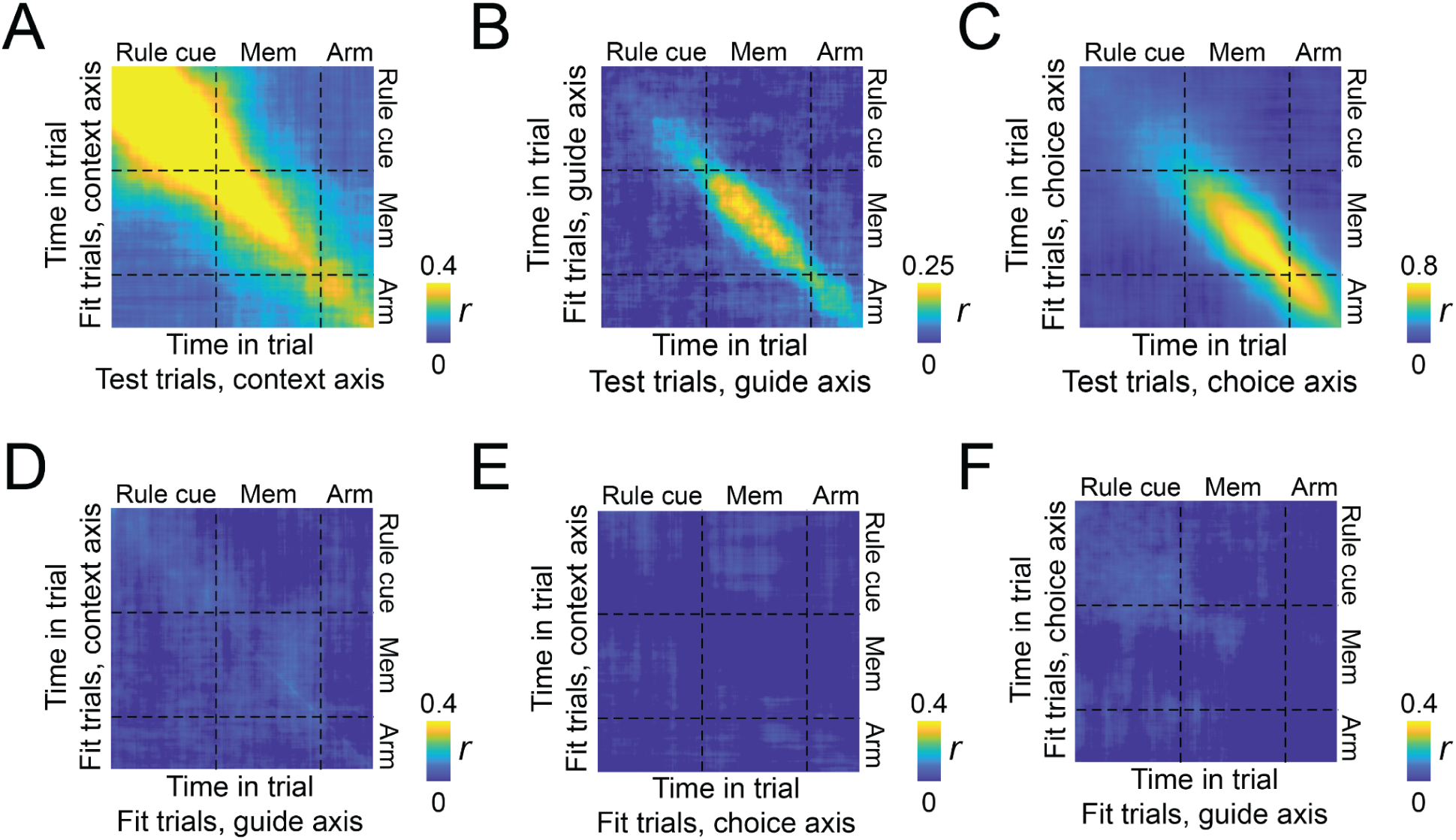
ProAnti task variables were represented along largely orthogonal axes of population activity. As expected for representations that modulate an underlying sequential activity structure, the axes for each variable tended to change over time. **(A-C)** Correlation of the (A) context, (B) guide, and (C) choice axes over time across two different halves of the laser-off trials. The context axis showed stability within the rule cue, but then changes over time in subsequent epochs. Both the guide and choice axes change over time during the memory period, which is when both guide and choice are represented. **(D-F)** Correlation between the (D) context vs. guide axes, (E) context vs. choice axes, and (F) guide vs. choice axes over time within the same set laser-off trials. At each timepoint, all three axes are largely orthogonal to each other.

**Figure S10.**
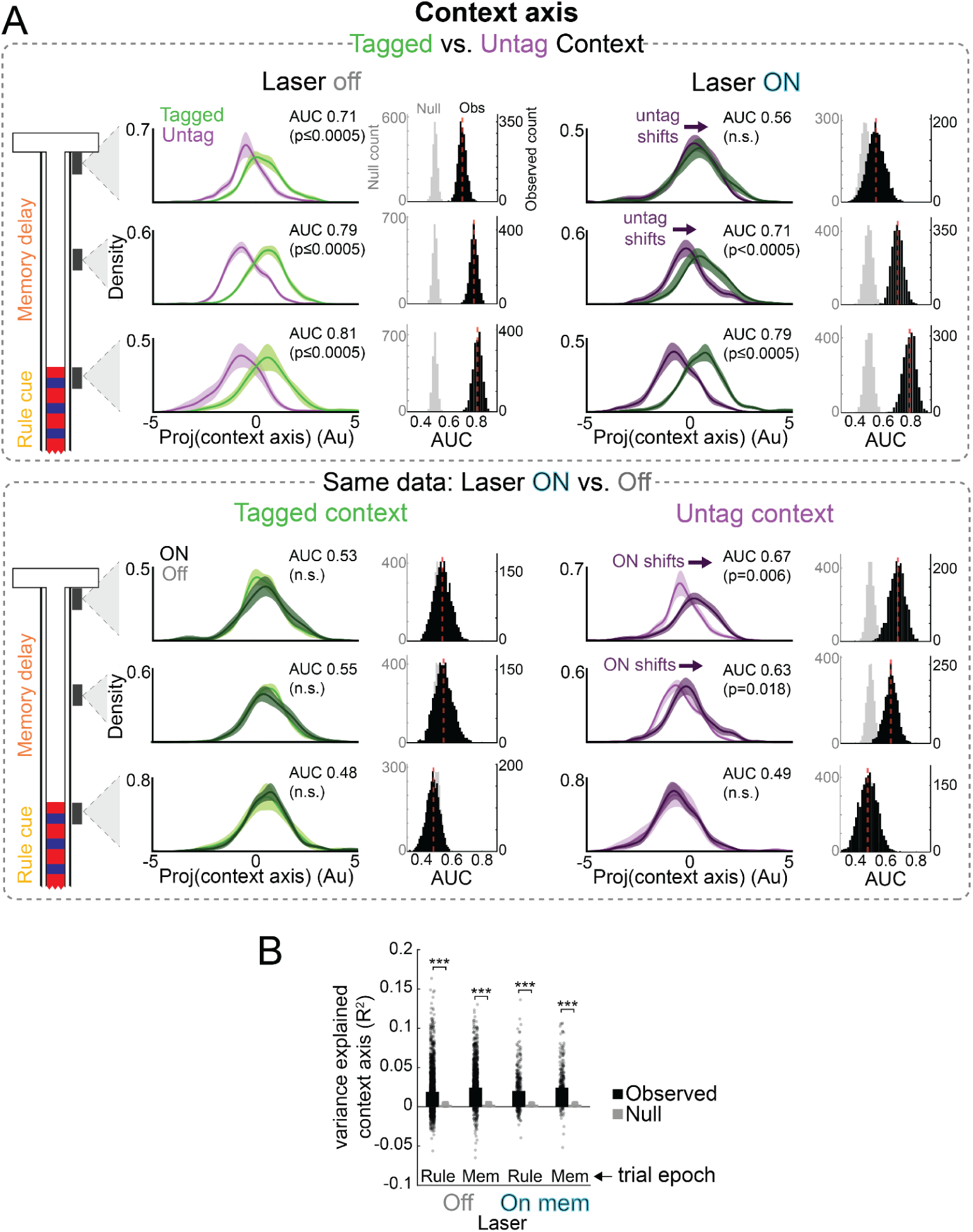
Memory delay HPC engram reactivation biases mPFC toward the tagged-context representation. **(A)** Distribution of population trajectories projected onto the context axis during laser-off and -on trials split by context (top), and tagged and untag context trials split by laser (bottom), during select task epoch windows (leftmost schematic) (bootstrap mean ± SEM; N=11 sessions). The laser was turned on during the memory delay. AUC for each bootstrap sample was compared in a pairwise fashion to a null AUC computed by shuffling trial labels (see Methods). P-values corrected for multiple comparisons. **(B)** Cross-validated variance explained (R^2^) by the context axis during memory delay reactivation trials (On) and laser-off trials (Off), when mice were in the rule cue zone (Rule) or memory delay (Mem) epoch. The context axis explained a greater fraction of variance across trials than expected by chance in each epoch during both laser-on and -off trials (observed vs. rotated null; linear mixed effects models, all t>14.8, ***p<10^-42^). There was no effect of the laser (2(trial type: off vs. on) x 2(epoch: rule vs mem) linear mixed-effects model; main effect laser: F_1,4586_=0.002, p=0.96; epoch x laser interaction; F_1,4586_=0.48, p=0.49). Dots denote individual trials.

**Figure S11.**
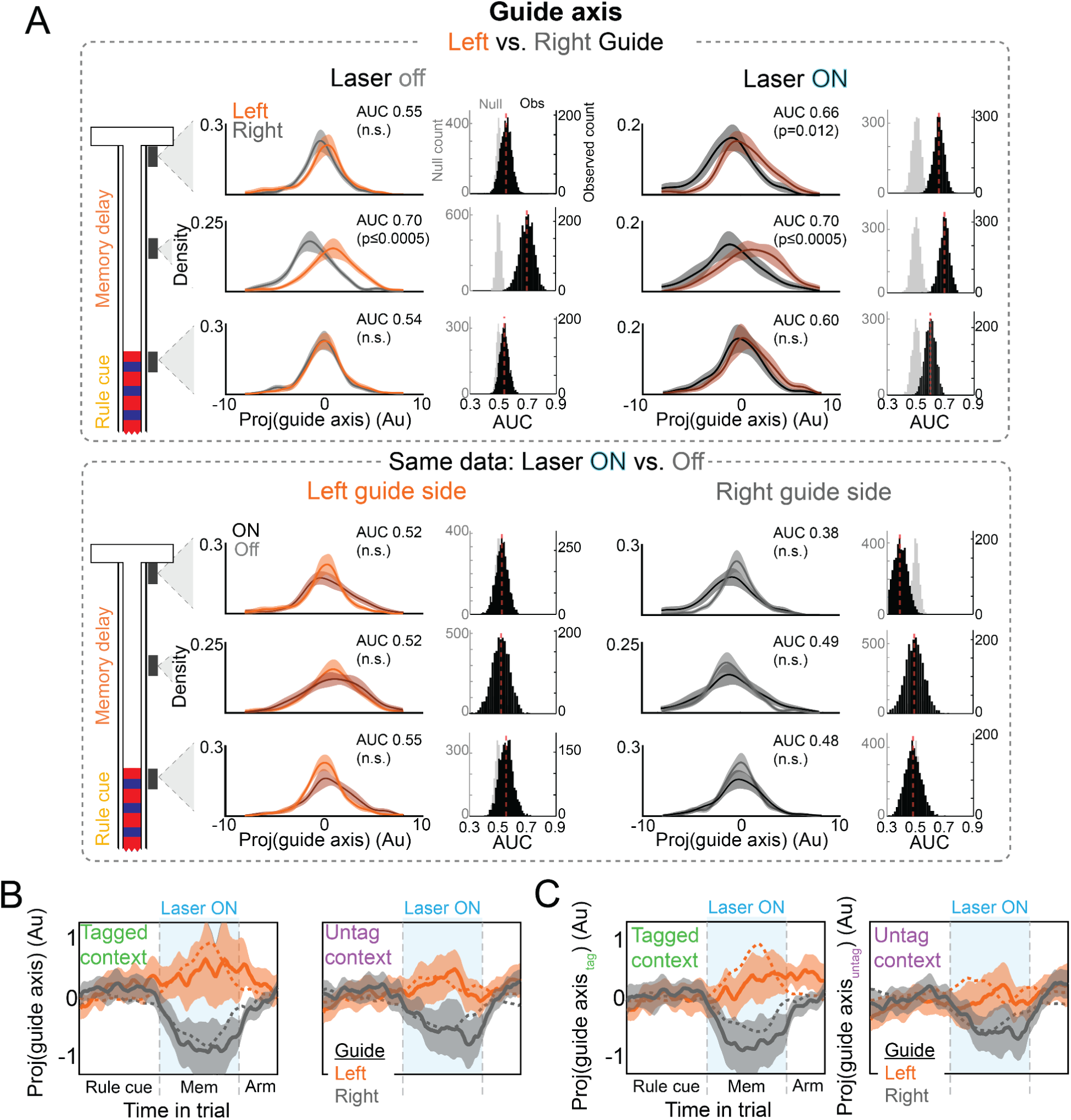
HPC engram reactivation does not affect mPFC guide side representations. **(A)** Same as Figure S10A, but for the guide axis. **(B)** Population trajectories projected onto the guide axis over time in a trial, separately in the tagged and untag contexts. No differences between laser-on and -off trials were observed in either context. **(C)** Same as (B) but with the guide axis computed separately in each context. No significant differences between laser-on and -off trials were observed in either context.

**Figure S12.**
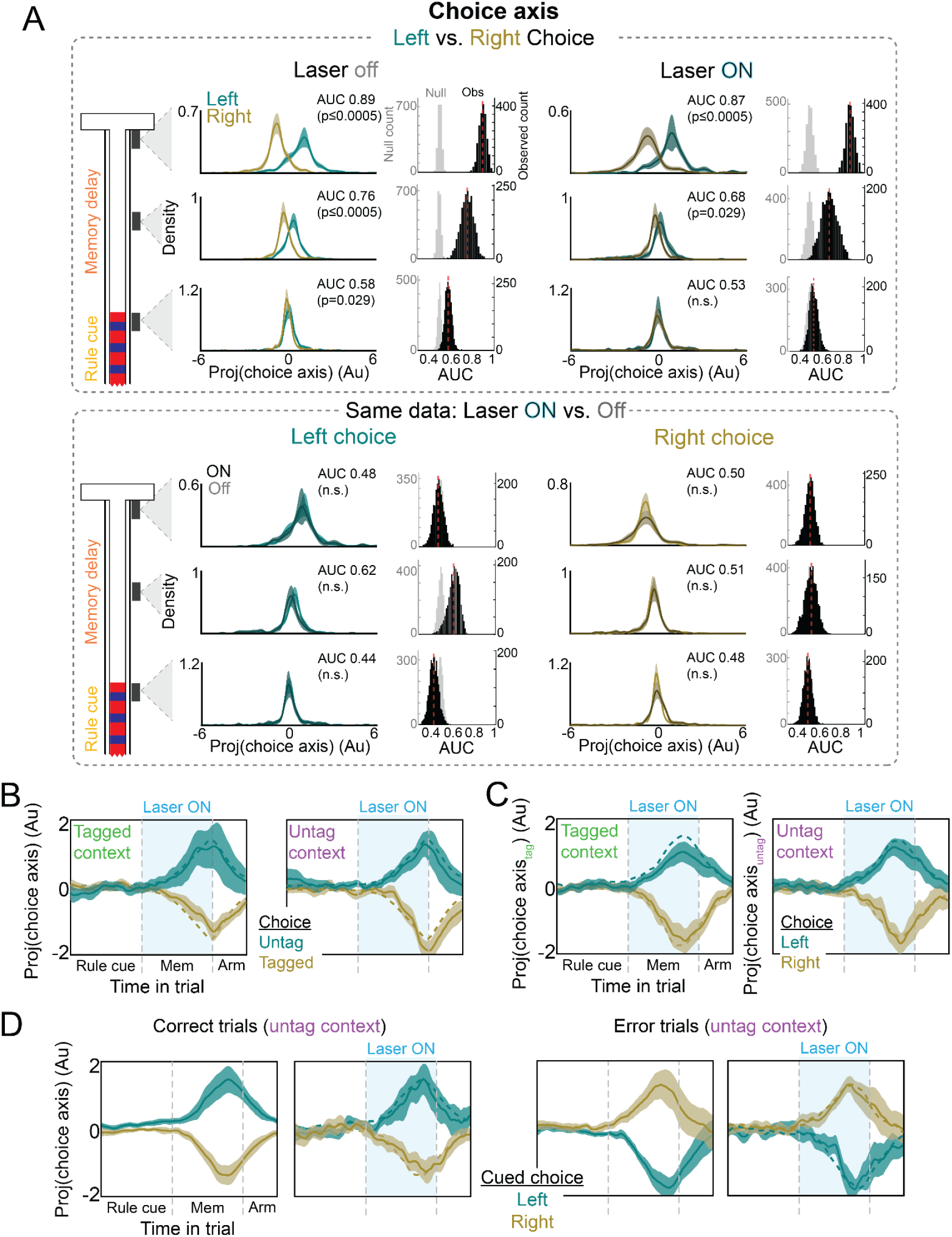
HPC engram reactivation does not affect mPFC choice representations. **(A)** Same as Fig. S10A but for the choice axis. **(B)** Population trajectories projected onto the choice axis over time in a trial, separately in the tagged and untag contexts. Projection values were computed here such that positive values correspond to choices in the untag direction, and negative values correspond to choices in the tagged direction. No significant differences between laser-on and -off trials were observed in either context. **(C)** Same as Fig. 3L but with the choice axis computed separately in each context. No significant differences between laser-on and -off trials were observed in either context. (**D**) Similar to Fig. 3H and 3L but during untag context correct (left) and error (right) trials. Population trajectories colors denote the experimentally cued correct choice, rather than the expressed choice. No significant differences between laser-on and -off trials were observed during either correct or error trials.

**Figure S13.**
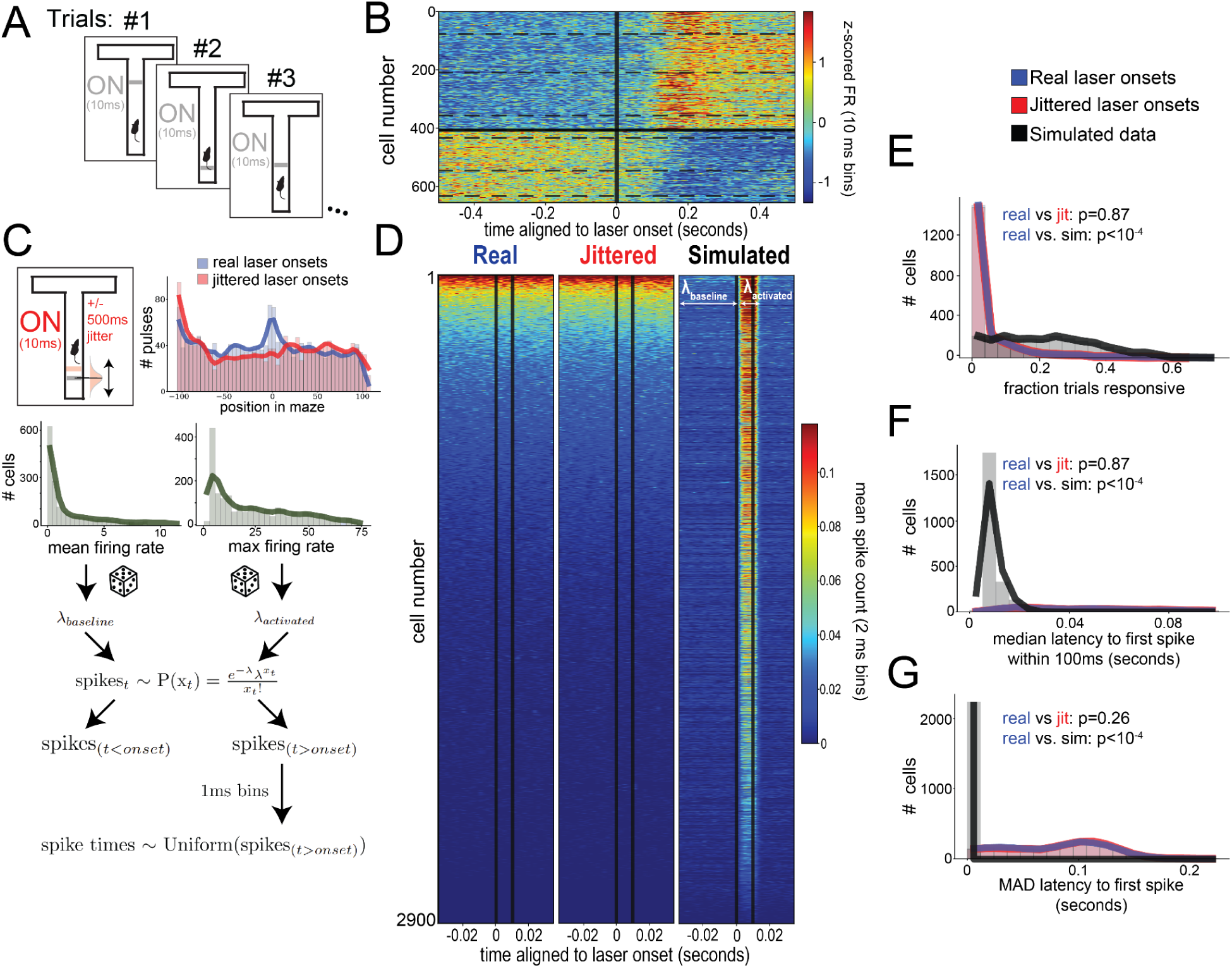
mPFC units were not directly activated by optogenetic stimulation. **(A)** Following the main-text reactivation experiments, separate mPFC recordings were collected during which a single 10ms pulse of light was delivered on every trial at a random location in the maze. **(B)** In the main-text experiments, individual mPFC units responded reliably to continuous 20 Hz stimulation with a latency of ∼100ms. Only significantly excited (top) or inhibited (bottom) cells (t-test, *p*<.01) are shown, evaluated across all trials and conditions. Dashed lines denote mouse identity, separately for excited and inhibited cells (N=4; cells are pooled from 2 sessions per mouse). **(C)** Jittered data was generated by randomly shifting the 10ms pulses in time with a gaussian centered around 0 with std=500ms (top left). Jittered onsets were distributed along the maze similarly to the actual onsets (top right). A third dataset was simulated by drawing firing rates from gaussians fit to either the mean or maximum firing rates across all cells during pre-laser baseline timepoints. These firing rates were then used to drive a Poisson process in 2ms bins, using means for bins prior to laser onset and maximums for bins following laser onset with a delay to accommodate ChR2 opening dynamics. Precise spike times for measuring latencies were drawn uniformly within each bin. **(D)** Mean spike counts in 2ms bins within a 70ms window aligned to laser onset. Vertical lines denote the 10ms pulse duration. **(E)** Distribution across cells within each dataset of the fraction of trials on which the cell responded to the laser pulse, defined as a mean firing rate during the pulse 1 standard deviation above the baseline mean. **(F)** Distribution of the median latency from laser onset to the first spike (ignoring spikes >100ms after the pulse). **(G)** Distribution of the median absolute deviation of the latency to all first spikes after the pulse. Across all metrics, firing rates aligned to actual laser onset times were statistically indistinguishable from firing rates aligned to randomly jittered laser onset times (KS tests).

**Figure S14.**
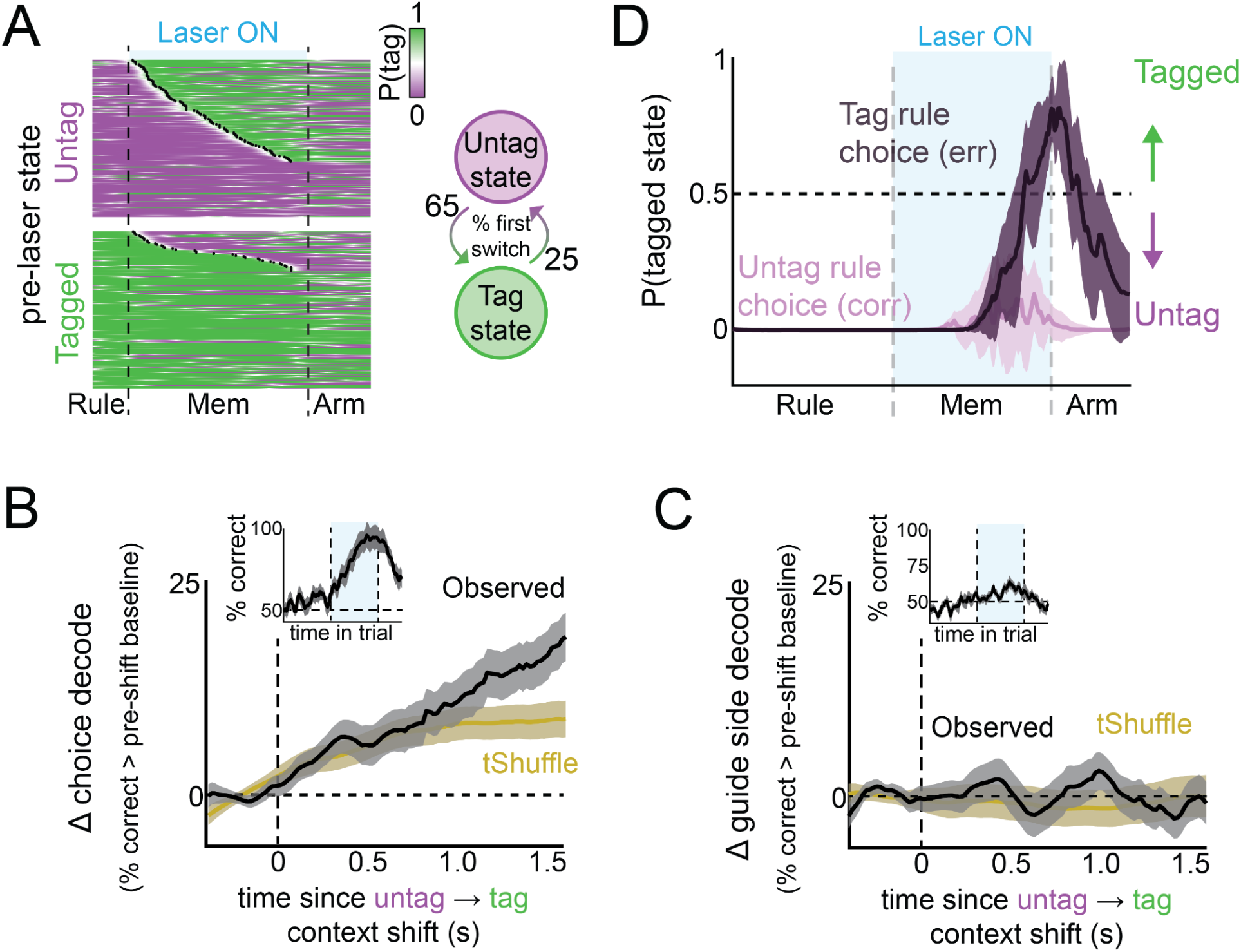
HPC engrams induce context state shifts at variable latencies, and choice representations emerge following the shift. **(A)** To identify when mPFC population dynamics transitioned between contextual states during HPC engram reactivation, we fit a two-state hidden Markov model (HMM) to laser-off context axis trajectories to estimate context-specific emission parameters, then applied this fixed-emission model to memory-delay reactivation trials to infer trial-by-trial state transition times. Left: HMM state probabilities over time during laser-on trials, grouped by model-defined pre-laser state, showing a preferential increase in transitions (black dots) from untag to tagged states during reactivation (χ^2^=54.9, p <10^-5^). On trials where a transition from the untagged to the tagged contextual state was detected, transition times were broadly distributed across the reactivation epoch, with a median latency of 849.8 ms following laser onset. Right: probability of remaining in the pre-laser state vs. switching for the first time. **(B)** Change in timepoint-by-timepoint LDA choice decoding accuracy from pre-laser baseline aligned to the trial-specific HMM-inferred context state transition time. The emergence of choice representations was time-locked to context state transitions, as compared to a temporally-shuffled state transition time null (tShuffle) (data vs. tShuffle: z = 3.59 over 1000-1600ms post-shift). Inset: LDA choice decoding accuracy relative to trial start during memory delay reactivation trials. Error bars denote hierarchical bootstrap mean ±1 SEM. **(C)** Same analysis as in (B) for guide decoding. Guide coding was unrelated to transition timing. **(D)** HMM-derived probability of being in the tagged context state over time during Untag context trials, split by expressed rule (Tag vs. Untag). Error bars denote bootstrap median ±1 SEM across trials.

**Figure S15.**
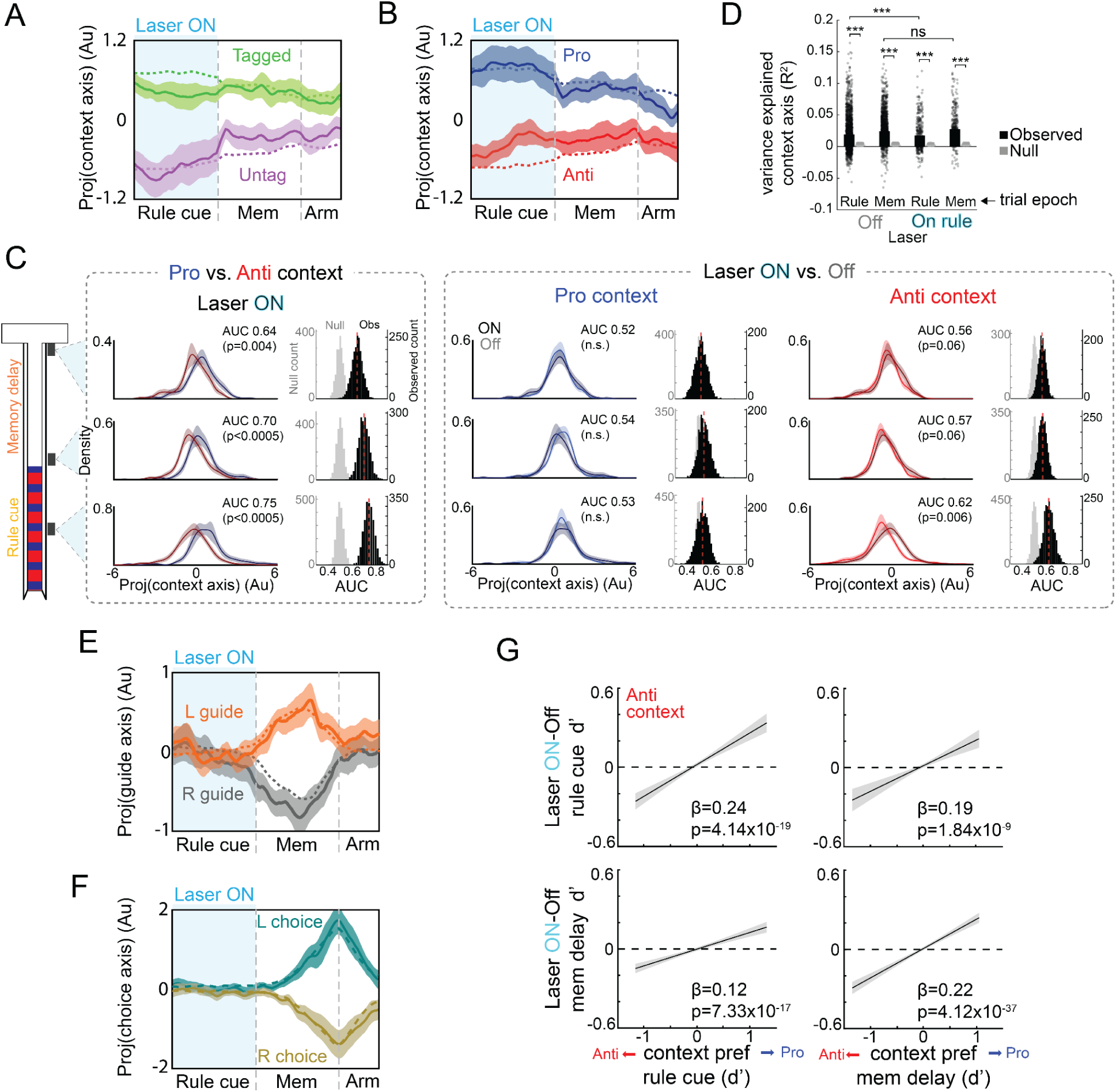
Rule-cue-only HPC engram reactivation causes ambiguous mPFC contextual representations. **(A)** Population trajectories projected onto the context axis during laser-off (dotted lines) and rule cue reactivation laser-on (shaded lines) trials for Tagged (green) and Untag (magenta) context trials. **(B)** Same data as in (A) but resorted according to the Pro (blue) and Anti (red) contexts, independent of tag identity. **(C)** Distribution of population trajectories projected onto the context axis split by context (left), and Pro and Anti context trials split by laser (right), during select task epoch windows (leftmost schematic) (bootstrap mean ± SEM; N=11 sessions). AUC for each bootstrap sample was compared in a pairwise fashion to a null AUC computed by shuffling trial labels (see Methods). There was a weak shift toward the Pro context trajectory during laser-on Anti context trials during the rule cue zone. P-values shown uncorrected. **(D)** Variance explained (R2) by the context axis during rule cue zone reactivation trials (On) and laser-off trials (Off), when mice were in the rule cue zone (Rule) or memory delay (Mem) epoch. The context axis was defined on independent trials from the R2 calculation. The context axis explained a greater fraction of variance across trials than expected by chance in all epochs for both laser-on and -off trials (observed vs. rotated null; linear mixed effects models, all t>14.8, p<10^-42^). However, the context axis accounted for less activity variance during laser-on than -off trials (2(trial type: off vs. on) x 2(epoch: rule vs mem) linear mixed-effects model; main effect laser: F_1,4586_=7.52, p=0.006), due to decreased variance explained in the rule cue zone (epoch x laser interaction; F_1,4658_=9.42, p=0.002). Dots denote individual trials. (**E-F**) Same as (A) but for (E) guide and (F) choice axes (bootstrap mean ± SEM; N=11 sessions). No significant differences between laser-on and -off trials were observed. **(G)** Mean laser modulation during the rule cue zone (top) and memory delay (bottom) vs. laser-off context discriminability (Anti vs. Pro-preferring) during the rule cue zone (left) and memory delay (right), pooled across all neurons (c.f. Figure 3C). Laser effects scaled with context tuning: anti-preferring neurons were suppressed and pro-preferring neurons excited (linear mixed model, shading denotes 95% CI).

**Figure S16.**
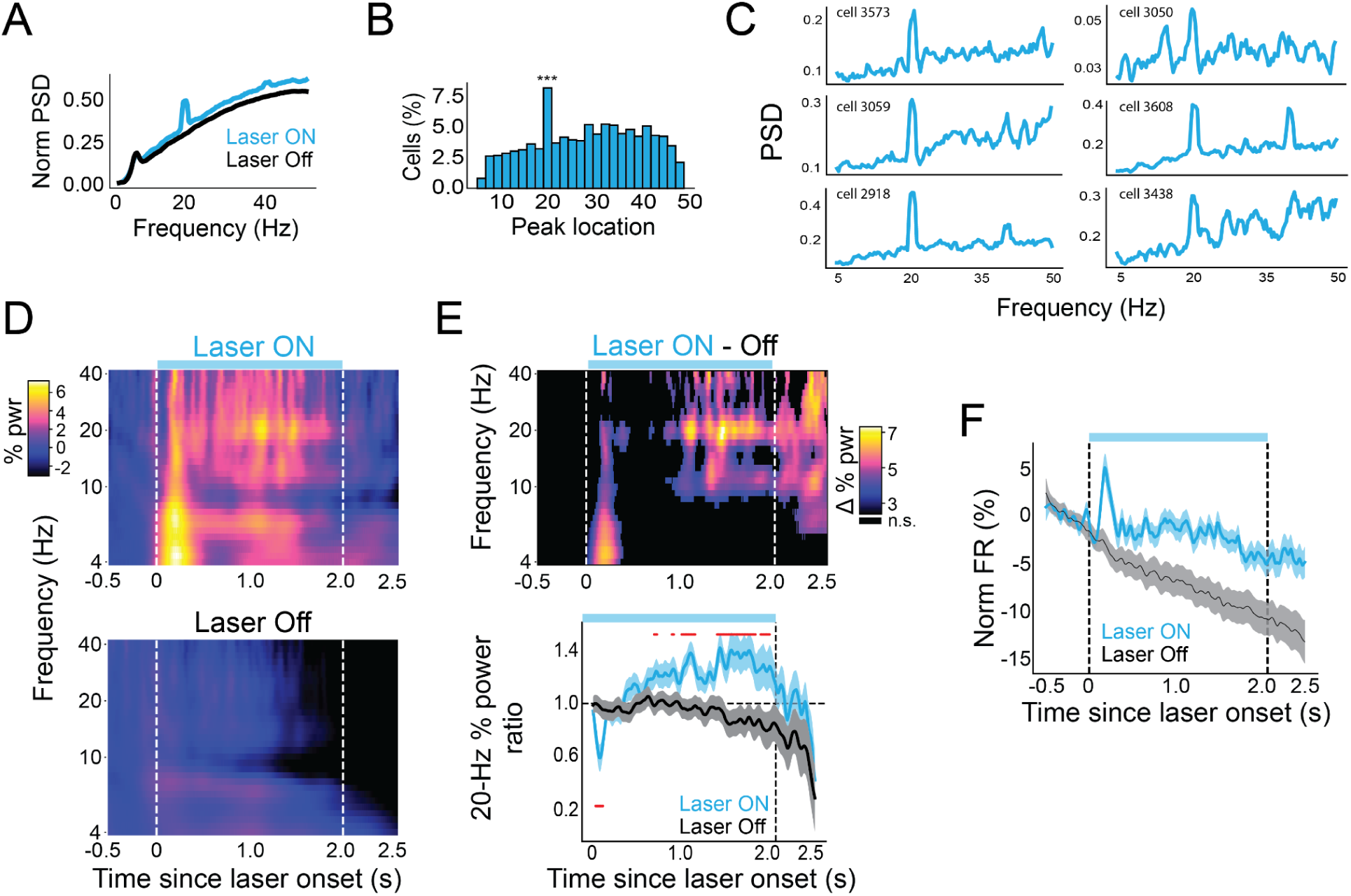
HPC engrams influence mPFC over synaptic and rapid network timescales. **(A)** Multitapering analysis averaged across all cells and memory zone laser-on trials (blue), or all laser-off trials (black). **(B)** Binned peaks from single cell power spectra averaged across trials (***binomial test, p_0_=0.083, p<10^-9^). **(C)** Illustrative example power spectra of single cells averaged across memory zone laser-on trials. Cells were selected randomly from those with peaks at 20 Hz. **(D)** Wavelet transform analysis aligned to memory zone onset for memory delay reactivation (upper left) or laser-off trials (bottom left), matched in choice statistics. Heat reflects percent change from a 500ms baseline averaged at each frequency. Vertical white lines denote laser onset and median laser offset time. **(E)** *Top*: Difference map between laser-on and matched laser-off trials (top), masked by significance (permutation test, FDR-corrected q<0.01). *Bottom*: The ratio of average percent change in the 20 Hz band to the average across all other frequencies. Red dots denote significant differences between laser-on and matched laser-off trials (permutation test, FDR-corrected q<0.01). **(F)** Percent change in firing rate (FR) relative to 500ms pre-laser baseline during laser-on and laser-off trials across the population (bootstrap mean ± 1 SD across cells). Individual cell FRs were smoothed with an acausal Gaussian filter (σ=20ms). There was a disproportionate spike in FR shortly (∼100ms) following laser onset compared to laser-off trials.

**Figure S17.**
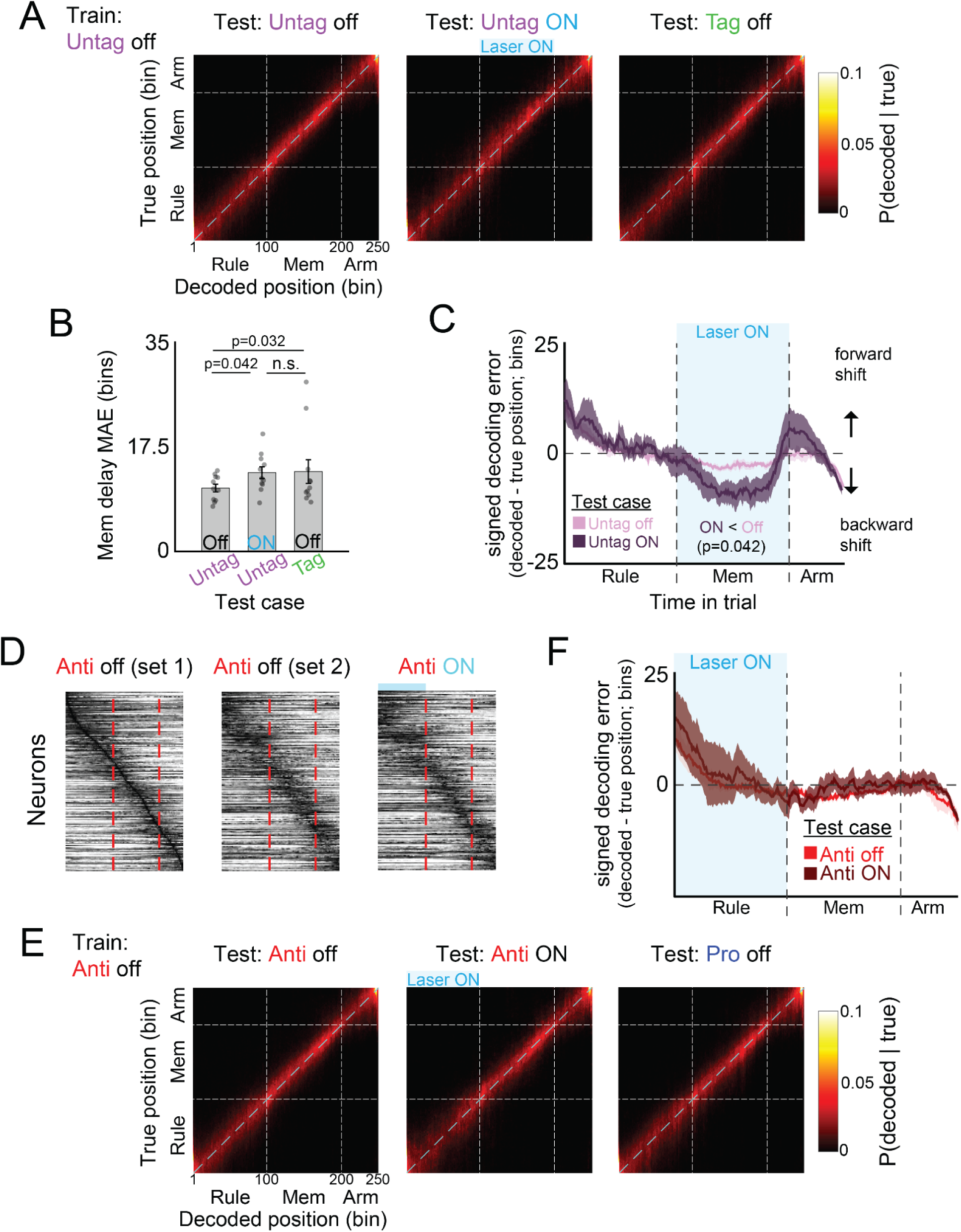
HPC engram reactivation does not disrupt mPFC sequential activity. **(A)** Time-resolved Bayesian decoding of spatiotemporal position using mPFC population activity. The decoder was trained on laser-off trials in the untagged context and tested on three conditions: laser-off untagged trials (independent test set), laser-on (memory delay reactivation) untagged trials, and laser-off tagged trials. Confusion matrices show true versus decoded position for each test condition. **(B)** Median absolute decoding error (MAE) during the memory delay for the three decoding conditions (untag off → untag off, untag off → untag memory delay laser-on, and untag off → tag off). There was a small but significant increase in MAE during laser-on compared to trials, comparable to the increase observed when decoding across contexts. Dots represent individual sessions and bars indicate session mean ± SEM. **(C)** Median signed decoding error over time during the trial, separately for laser-off untagged trials (independent test set) and laser-on (memory delay reactivation) untagged trials. The decoder predicted a small backward shift in decoded position during HPC stimulation (averaged over the memory delay) during laser-on relative to laser-off trials. In (B-C) statistical comparisons performed using Wilcoxon signed-rank tests. **(D)** Anti context activity traces sorted by peak firing time during an example laser-on–trial-matched subset of laser-off trials (set 1), and during rule-cue-only HPC reactivation trials. Rows are max-normalized per condition. **(E)** Same as (A) but for a decoder trained on laser-off trials in the Anti context and tested during: laser-off Anti trials (independent test set), laser-on (rule cue reactivation) Anti trials, and laser-off Pro trials. **(F)** Same as (C) separately for laser-off Anti trials (independent test set) and laser-on (rule cue zone reactivation) Anti trials. No significant difference between laser-on and laser-off trials.

